# Uncertainpy: A Python toolbox for uncertainty quantification and sensitivity analysis in computational neuroscience

**DOI:** 10.1101/274779

**Authors:** Simen Tennøe, Geir Halnes, Gaute T. Einevoll

**Affiliations:** Centre for Integrative Neuroplasticity, University of Oslo, Oslo, Norway; Department of Informatics, University of Oslo, Oslo, Norway; Faculty of Science and Technology, Norwegian University of Life Sciences, Ås, Norway; Department of Physics, University of Oslo, Oslo, Norway

**Keywords:** uncertainty quantification, sensitivity analysis, features, polynomial chaos, quasi-Monte Carlo methods, stochastic modeling, computational modeling, Python

## Abstract

Computational models in neuroscience typically contain many parameters that are poorly constrained by experimental data. Uncertainty quantification and sensitivity analysis provide rigorous procedures to quantify how the model output depends on this parameter uncertainty. Unfortunately, the application of such methods is not yet standard within the field of neuroscience.

Here we present Uncertainpy, an open-source Python toolbox, tailored to perform uncertainty quantification and sensitivity analysis of neuroscience models. Uncertainpy aims to make it easy and quick to get started with uncertainty analysis, without any need for detailed prior knowledge. The toolbox allows uncertainty quantification and sensitivity analysis to be performed on already existing models without needing to modify the model equations or model implementation. Uncertainpy bases its analysis on polynomial chaos expansions, which are more efficient than the more standard Monte-Carlo based approaches.

Uncertainpy is tailored for neuroscience applications by its built-in capability for calculating characteristic features in the model output. The toolbox does not merely perform a point-to-point comparison of the “raw” model output (e.g. membrane voltage traces), but can also calculate the uncertainty and sensitivity of salient model response features such as spike timing, action potential width, mean interspike interval, and other features relevant for various neural and neural network models. Uncertainpy comes with several common models and features built in, and including custom models and new features is easy.

The aim of the current paper is to present Uncertainpy for the neuroscience community in a user-oriented manner. To demonstrate its broad applicability, we perform an uncertainty quantification and sensitivity analysis on three case studies relevant for neuroscience: the original Hodgkin-Huxley point-neuron model for action potential generation, a multi-compartmental model of a thalamic interneuron implemented in the NEURON simulator, and a sparsely connected recurrent network model implemented in the NEST simulator.

**SIGNIFICANCE STATEMENT:** A major challenge in computational neuroscience is to specify the often large number of parameters that define the neuron and neural network models. Many of these parameters have an inherent variability, and some may even be actively regulated and change with time. It is important to know how the uncertainty in model parameters affects the model predictions. To address this need we here present Uncertainpy, an open-source Python toolbox tailored to perform uncertainty quantification and sensitivity analysis of neuroscience models.

## 1 INTRODUCTION

Computational modeling has become a useful tool for examining various phenomena in biology in general (Brodland, 2015) and neuroscience in particular (Koch and Segev, 1998; Dayan and Abbott, 2001; Sterratt et al., 2011). The field of neuroscience has seen the development of ever more complex models, and models now exist for large networks of biophysically detailed neurons (Izhikevich and Edelman, 2008; Merolla et al., 2014; Markram et al., 2015).

Computational models typically contain a number of parameters that for various reasons are uncertain. A typical example of an uncertain parameter in a neural model can be the conductance *g*_*x*_ of a fully open ion channel of a specific type *x*. Despite the parameter uncertainty, it is common practice to construct models that are deterministic in the sense that single numerical values are assigned to each parameter.

Uncertainty quantification is a means to quantify the uncertainty in the model output that arise from uncertainty in the model parameters. Instead of using fixed model parameters as in a deterministic model (as illustrated in Figure 1A), one assigns a distribution of possible values to each model parameter. The uncertainties in the model parameters are then propagated through the model and give rise to a distribution in the model output (as illustrated in Figure 1B). An uncertainty quantification can thus be seen as a transformation from a deterministic model to a stochastic model.

**Figure 1.**
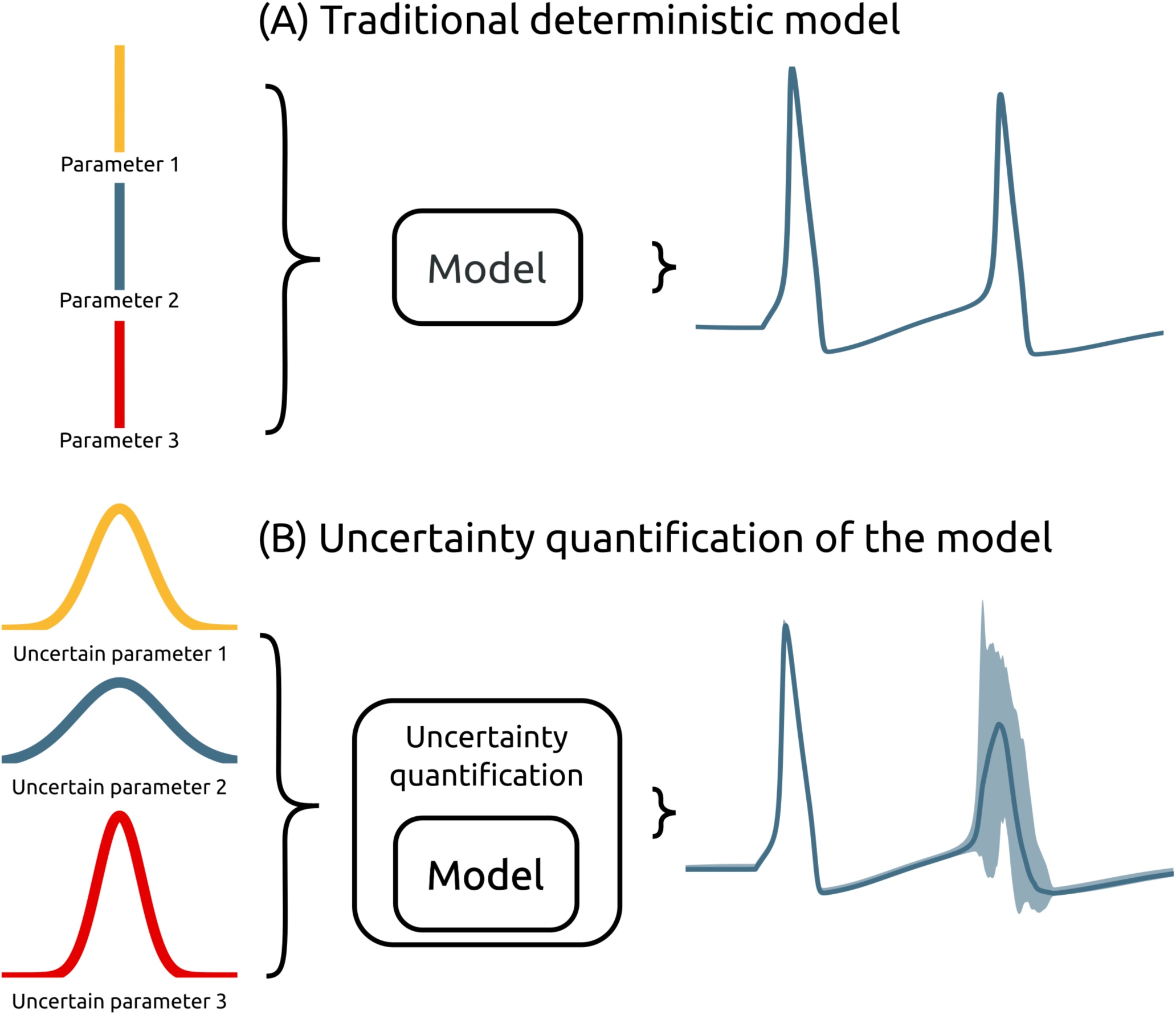
Illustration of an uncertainty quantification of a deterministic model. (A) A traditional deterministic model where each input parameter has a fixed value, and we get one output of the model (grey). (B) An uncertainty quantification of the model makes the parameters of the model have distributions, and the output of the model becomes a range of possible values (light grey).

Sensitivity analysis is tightly linked to uncertainty quantification, and is the process of quantifying how much of the output uncertainty each parameter is responsible for (Saltelli, 2002). In addition to providing insight into how different aspects of the model affects its overall properties, knowing the sensitivity of the model to each parameter can be used in guiding experimental analysis, model reduction and parameter estimation (Degenring et al., 2004; Zi, 2011; Snowden et al., 2017).

The uncertainty in a model parameter may have many origins. It may be due to (i) measurement uncertainty or (ii) lack of experimental techniques that enables the parameter to be measured. The uncertainty can also be due to an inherent biological variability, meaning the value of a parameter can vary (iii) between neurons of the same species (Edelman and Gally, 2001; Hay et al., 2013), or (iv) dynamically within a single neuron due to plasticity or homeostatic mechanisms (Marder and Goaillard, 2006). Additionally, some models include parameters that are (v) phenomenological abstractions, and therefore do not represent directly measurable physical entities. They might for example represent the combined effect of several physical processes.

A common way to avoid addressing the uncertainty in measured parameters is to use the mean of several experimental measurements. This can be problematic since the underlying distribution of a set of parameters can be poorly characterized by the mean and variance of each parameter (Golowasch et al., 2002). Additionally, during model construction, a subset of the uncertain parameters are commonly treated as *free parameters*. This means the parameters are tuned by the modeller to values that makes the model output match a set of experimental constraints. An example is fitting an ion-channel conductance *g*_*x*_ so the membrane potential of a neuron model reproduces an experimentally measured voltage trace. Historically, such parameter tuning was often done manually by trial and error, but a variety of automated parameter fitting algorithms have gradually taken over (Bhalla and Bower, 1993; Vanier and Bower, 1999; Druckmann et al., 2007; Van Geit et al., 2007, 2008; Taylor et al., 2009; Hay et al., 2011; Svensson et al., 2012; Bahl et al., 2012; Friedrich et al., 2014; Pozzorini et al., 2015; Van Geit et al., 2016; Mäki-Marttunen et al., 2018). Whatever method used, the tuning procedure does not guarantee a unique solution for the *correct* parameter set, since it often is the case that a wide range of different parameter combinations give rise to similar model behavior (Bhalla and Bower, 1993; Beer et al., 1999; Goldman et al., 2001; Golowasch et al., 2002; Prinz et al., 2004; Tobin, 2006; Schulz et al., 2007; Halnes et al., 2007; Taylor et al., 2009; Marder and Taylor, 2011).

Given that most neuroscience models contain a variety of uncertain parameters, the need for systematic approaches to quantify what confidence we can have in the model output is pressing. The importance of uncertainty quantification and sensitivity analysis of computational models is well known in a wide variety of fields (Leamer, 1985; Beck, 1987; Turanyi and Turányi, 1990; Oberkampf et al., 2002; Wood-Schultz, 2011; Marino et al., 2008; Najm, 2009; Rossa et al., 2011; Yildirim and Karniadakis, 2015; Wang and Sheen, 2015). Due the prevalence of inherent variability in the parameters of biological systems, uncertainty quantification and sensitivity analysis is at least as important in neuroscience.

Unfortunately, a generally accepted practice for uncertainty quantification and sensitivity analysis does not currently exist within the field of neuroscience, and models are commonly presented without including any form of uncertainty quantification or sensitivity analysis. When an effort is made in that direction, it is still common to use rather simple, so called One-At-A-Time methods, where one examines how much the model output changes when varying one parameter at a time (see e.g., De Schutter and Bower (1994); Blot and Barbour (2014); Kuchibhotla et al. (2017)). Such approaches do not account for potential dependencies between the parameters, and thereby miss correlations within the often multi- dimensional parameter space (Borgonovo and Plischke, 2016). Other methods that have been applied are local methods, which are multi-dimensional, but confined to exploring small perturbations surrounding a single point in the parameter space (see e.g., Gutenkunst et al. (2007); Blomquist et al. (2009); O’Donnell et al. (2017)). Such methods can thus not explore the effect of arbitrarily broad uncertainty distributions for the parameters.

Methods for uncertainty quantification and sensitivity analysis that take the entire parameter space into account are often called global methods (Borgonovo and Plischke, 2016; Babtie and Stumpf, 2017). Global methods are only occasionally used within the field of neuroscience (see e.g., Torres Valderrama et al. (2015); Halnes et al. (2009)). The most well known of the global methods are (quasi-)Monte Carlo methods, which rely on randomly sampling the parameter distributions, and then calculate statistics from the following model outputs. The problem with (quasi-)Monte Carlo methods is that they are computationally very demanding, particularly for large and complex models. A means to obtain similar results in a much more efficient way, is provided by the recent mathematical framework of polynomial chaos expansions (Xiu and Hesthaven, 2005). Polynomial chaos expansions are used to approximate the model with a polynomial (as a surrogate model), on which the uncertainty and sensitivity analysis can be performed much more efficiently.

To lower the threshold for neuroscientists to perform uncertainty quantification and sensitivity analysis we have created Uncertainpy^1^, an open-source Python toolbox for efficient uncertainty quantification and sensitivity analysis. Uncertainpy aims to make it easy and quick to get started with uncertainty quantifications and sensitivity analysis, just a few lines of Python code are needed, without any need for detailed prior knowledge of uncertainty analysis. Uncertainpy implements both quasi-Monte Carlo methods and polynomial chaos expansions. The toolbox is model independent and treats the model as a “black box”, meaning that uncertainty quantification can be performed on already existing models without needing to modify the model equations or model implementation. We hope Uncertainpy can help to enable uncertainty quantification and sensitivity analysis in neuroscience.

Whereas its statistical methods are generally applicable, Uncertainpy is tailored for neuroscience applications by having an built-in capability for recognizing characteristic features in the model output. This means Uncertainpy does not merely perform a point-to-point comparison of the “raw” model output (e.g., a voltage trace). When applicable, Uncertainpy also recognizes and calculates the uncertainty in model response features, for example the spike timing and action potential shape for neural models, and measurements of spiking neural network activity, such as firing rates and interspike intervals, for neural networks.

To present Uncertainpy, we start this paper with an overview of the theory behind uncertainty quantification and sensitivity analysis in Section 2, with a focus on (quasi-)Monte Carlo methods and polynomial chaos expansions. In Section 3 we explain how to use Uncertainpy, and give details on how the uncertainty quantification is implemented. In Section 4 we illustrate the use of Uncertainpy by showing four different case studies where we perform uncertainty analysis of: (i) a cooling coffee-cup model (Newton’s law of cooling) to start with a simple use case, (ii) the original Hodgkin-Huxley point-neuron model for action potential generation, (iii) a comprehensive multi-compartmental model of a thalamic interneuron, and (iv) a sparsely connected recurrent network model. We end with a discussion and some future prospects in Section 5.

## 2 THEORY ON UNCERTAINTY QUANTIFICATION AND SENSITIVITY ANALYSIS

Uncertainty quantification and sensitivity analysis provide rigorous procedures to analyse and characterize the effects of parameter uncertainty on the output of a model. The methods for uncertainty quantification and sensitivity analysis can be divided into global and local methods. Local methods examine how the model output changes with small perturbations around a fixed point in the parameter space. Global methods, on the other hand, consider the entire parameter space. Global methods can therefore identify complex dependencies between the model parameters in terms of how they affect the model output.

The global methods can be further divided into intrusive and non-intrusive methods. Intrusive methods require changes to the underlying model equations, and are often challenging to implement. Models in neuroscience are often created with the use of advanced simulators such as NEST (Eppler et al., 2015) and NEURON (Hines and Carnevale, 1997). Modifying the underlying equations of models using these simulators is a complicated task best avoided. Non-intrusive methods, on the other hand, consider the model as a black box, and can be applied to any model without needing to modify the model equations or model implementation. Global, non-intrusive methods are therefore the methods of choice in Uncertainpy. The uncertainty calculations in Uncertainpy is based on the Python package *Chaospy* (Feinberg and Langtangen, 2015), which provides global non-intrusive methods for uncertainty quantification and sensitivity analysis.

In this section we go through the theory behind the methods for uncertainty quantification and sensitivity analysis used in Uncertainpy. We start by introducing the notation used in this paper (Section 2.1). Next, we introduce the statistical measurements for uncertainty quantification (Section 2.2) and sensitivity analysis (Section 2.3). Further, we give an introduction to (quasi-)Monte Carlo methods (Section 2.4) and polynomial chaos expansions (Section 2.5), the two methods used to perform the uncertainty analysis in Uncertainpy. We next explain how Uncertainpy handle cases with statistically dependent model parameters (Section 2.6). Finally, we explain the concept and benefits of performing a feature-based analysis (Section 2.7). We note that detailed insight into the theory of uncertainty quantification and sensitivity analysis is not a prerequisite for *using* Uncertainpy, so the more practical oriented reader may chose to skip this section, and go directly to the user guide in Section 3.

### 2.1 Problem definition

Consider a model *U* that depends on space ***x*** and time *t*, has *D* uncertain input parameters ***Q*** = [*Q*_1_*, Q*_2_*, …, Q*_*D*_], and gives the output *Y*:

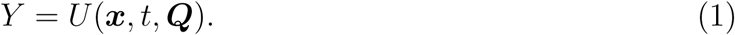

The output *Y* can have any value within the output space Ω_*Y*_ and has an unknown probability density function *ρ*_*Y*_. The goal of an uncertainty quantification is to describe the unknown *ρ*_*Y*_ through statistical metrics. We are only interested in the input and output of the model, and we ignore all details on how the model works. The model *U* is thus considered a black box, and may represent any model, for example a spiking neuron model that returns a voltage trace, or a neural network model that returns a spike train.

We assume the model includes uncertain parameters that can be described by a multivariate probability density function *ρ***_*Q*_**. Examples of parameters that can be uncertain in neuroscience are the conductance of a single ion channel, or the synaptic weight between two types of neurons in a neural network. If the uncertain parameters are independent, the multivariate probability density function *ρ***_*Q*_** can be given as separate univariate probability density functions *ρ*_*Q*_*i*, one for each uncertain parameter *Q*_*i*_. The joint multivariate probability density function for the independent uncertain parameters is then:

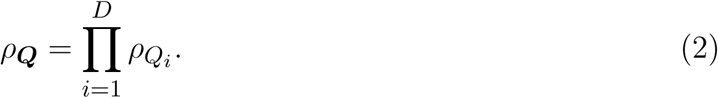

In cases where the uncertain input parameters are dependent, the multivariate probability density function *ρ***_*Q*_** must be defined directly. We assume the probability density functions are known, and are not here concerned with how they are determined. They may be the product of a series of measurements, a parameter estimation, or educated guesses made by experts.

### 2.2 Uncertainty quantification

As mentioned, the goal of an uncertainty quantification is to describe the unknown distribution of the model output *ρ*_*Y*_ through statistical metrics. The two most common statistical metrics used in this context are the mean 𝔼 (also called the expectation value) and the variance V. The mean is defined as:

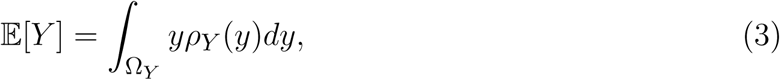

and tells us the expected value of the model output *Y*. The variance is defined as:

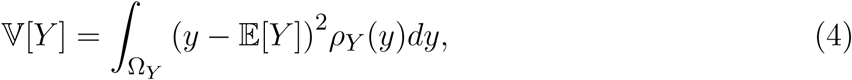

and tells us how much the output varies around the mean.

Another useful metric is the (100 *x*)-th percentile *P*_*x*_ of *Y*, which defines a value below which 100 *x* percent of the simulation outputs are located. For example, 5% of the simulations of a model will give an output lower than the 5-th percentile. The (100 *x*)-th percentile is defined as:

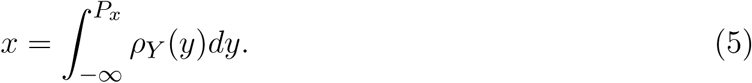

We can combine two percentiles to create a prediction interval *I*_*x*_, which is a range of values such that a 100 · *x* percentage of the outputs *Y* occur within this range:

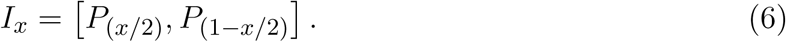

The 90% prediction interval gives us the interval within 90% of the *Y* outcomes occur, which also means that 5% of the outcomes are above and 5% below this interval.

### 2.3 Sensitivity analysis

Sensitivity analysis quantifies how much of the uncertainty in the model output each uncertain parameter is responsible for. It is the computational equivalent of analysis of variance (ANOVA) performed on experimental data (Archer et al., 1997). For a review of different sensitivity analysis methods, see Hamby (1994); Borgonovo and Plischke (2016). Several different sensitivity measures exist, but Uncertainpy uses the commonly used Sobol sensitivity indices (Sobol, 1990). The Sobol sensitivity indices quantify how much of the variance in the model output each uncertain parameter is responsible for. If a parameter has a low sensitivity index, variations of this parameter results in comparatively small variations in the final model output. On the other hand, if a parameter has a high sensitivity index, a change in this parameter leads to a large change in the model output.

A sensitivity analysis provides a better understanding of the relationship between the parameters and output of a model. This can be useful in a model reduction context. For example, a parameter with a low sensitivity index can essentially be set to any fixed value (within the explored distribution), without affecting the variance of the model much. In some cases, such an analysis can justify leaving out entire mechanisms from a model. For example, if a single neuron model is insensitive to the conductance of a given ion channel *g*_*x*_, this ion channel could possibly be removed from the model without changing the model behavior much. Additionally, a model-based sensitivity analysis can guide the experimental focus, so that special care is taken to obtain accurate measures of parameters with high sensitivity indices, while more crude measures are acceptable for parameters with low sensitivity indices.

There exist several types of Sobol indices. The first-order Sobol sensitivity index *S* measures the direct effect each parameter has on the variance of the model:

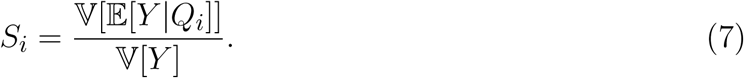

Here, 𝔼 [*Y Q*_*i*_] denotes the expected value of the output *Y* when parameter *Q*_*i*_ is fixed. The first-order Sobol sensitivity index tells us the expected reduction in the variance of the model when we fix parameter *Q*_*i*_. The sum of the first-order Sobol sensitivity indices can not exceed one (Glen and Isaacs, 2012).

Higher order sobol indices exist, and give the sensitivity due interactions between a parameter *Q*_*i*_ and various other parameters. It is customary to only calculate the first and total-order indices (Saltelli et al., 2010). The total Sobol sensitivity index *S*_*T* *i*_ includes the sensitivity of both the first-order effects, as well as the sensitivity due to interactions (covariance) between a given parameter *Q*_*i*_ and all other parameters (Homma and Saltelli, 1996). It is defined as:

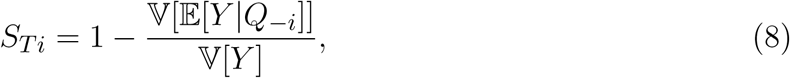

where *Q* _–*i*_ denotes all uncertain parameters except *Q*_*i*_. The sum of the total Sobol sensitivity indices is equal to or greater than one (Glen and Isaacs, 2012). If no higher order interactions are present, the sum of both the first and total-order sobol indices are equal to one.

We might want to compare Sobol indices across different features (introduced in Section 2.7). This can be problematic when we have features with different number of output dimensions. In the case of a zero-dimensional output the Sobol indices is a single number, while for a one-dimensional output we get Sobol indices for each point in time. To better be able to compare the Sobol indices across such features, we therefore calculate the normalized sum of both the first-order Sobol indices *Ŝ*, and the total-order Sobol indices *Ŝ*_*T*_

### 2.4 (Quasi-)Monte Carlo methods

A typical way to obtain the statistical metrics mentioned above is to use (quasi-)Monte Carlo methods. We give a brief overview of these methods here, for more comprehensive reviews see Lemieux (2009); Rubinstein and Kroese (2016).

The general idea behind the standard Monte Carlo method is quite simple. A set of parameters is randomly drawn from the joint multivariate probability density function *ρ***_*Q*_** of the parameters. The model is then evaluated for the sampled parameter set. This process is repeated thousands of times, and statistical metrics such as the mean and variance are computed for the resulting series of model outputs. The problem with the standard Monte Carlo method is that a very high number of model evaluations is required to get reliable statistics. If the model is computationally expensive, the Monte Carlo method may require insurmountable computer power.

Quasi-Monte Carlo methods improve upon the standard Monte Carlo method by using variance-reduction techniques to reduce the number of model evaluations needed. These methods are based on increasing the coverage of the sampled parameter space by distributing the samples more evenly. Fewer samples are then required to get a given accuracy. Instead of randomly selecting parameters from *ρ***_*Q*_**, the samples are selected using a low-discrepancy sequence such as the Hammersley sequence (Hammersley, 1960), which is used in Uncertainpy. Quasi-Monte Carlo methods are faster than the Monte Carlo method, as long as the number of uncertain parameters is sufficiently small (Lemieux, 2009).

Uncertainpy allows quasi-Monte Carlo methods to be used to compute the statistical metrics. When this option is chosen, the metrics are computed as follows. With *N* model evaluations, which gives the results ***Y*** = [*Y*_1_*, Y*_2_*, …, Y*_*N*_], the mean is given by

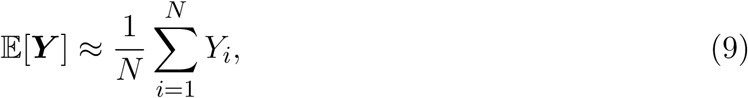

and the variance by

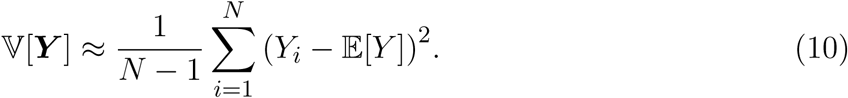

Prediction intervals are found by sorting the model evaluations ***Y*** in an increasing order, and then finding the (100 *x*/2)-th and (100 · (1 – *x*/2))-th percentiles. The sensitivity analysis in Uncertainpy is based on polynomial chaos expansions (see below), and Uncertainpy does currently not support calculation of Sobol indices from (quasi-)Monte Carlo methods, although methods for this are available in the literature (Saltelli et al., 2010).

### 2.5 Polynomial chaos expansions

A recent mathematical framework for estimating uncertainty is that of polynomial chaos expansions (Xiu and Hesthaven, 2005). Polynomial chaos expansions can be seen as a subset of polynomial approximation methods. For a review of polynomial chaos expansions see Xiu (2010). Polynomial chaos expansions are much faster than (quasi-)Monte Carlo methods as long as the number of uncertain parameters is relatively low, typically smaller than about twenty (Crestaux et al., 2009). This is the case for many neuroscience models, and even for models with a higher number of uncertain parameters, the analysis can be performed for selected subsets of the parameters.

The general idea behind polynomial chaos expansions is to approximate the model *U* with a polynomial expansion *Û*:

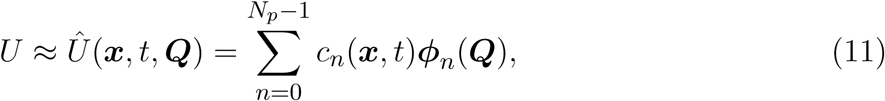

where ***ϕ****n* denote polynomials, and *c*_*n*_ denote expansion coefficients. The number of expansion factors *N*_*p*_ is given by

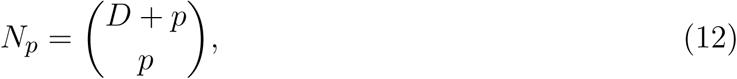

where *p* is the polynomial order. The polynomials *ϕ*_*n*_(***Q***) are chosen so they are orthogonal with respect to the probability density function *ρ***_*Q*_**, which ensures useful statistical properties.

When creating the polynomial chaos expansion, the first step is to find the orthogonal polynomials ***ϕ****n*, which in Uncertainpy is done using the so called three-term recurrence relation (Xiu, 2010). The next step is to estimate the expansion coefficients *c*_*n*_. The non-intrusive methods for doing this can be divided into two classes, point-collocation methods and pseudo-spectral projection methods, both of which are implemented in Uncertainpy.

Point collocation is the default method used in Uncertainpy. This method is based on demanding that the polynomial approximation is equal to the model output evaluated at a set of collocation nodes drawn from the joint probability density function *ρ***_*Q*_**. This demand results in a set of linear equations for the polynomial coefficients *c*_*n*_, which can be solved by the use of regression methods. The regression method used in Uncertainpy is Tikhonov regularization (Rifkin and Lippert, 2007).

Pseudo-spectral projection methods are based on least squares minimization in the orthogonal polynomial space, and finds the expansion coefficients *c*_*n*_ through numerical integration. The integration uses a quadrature scheme with weights and nodes, and the model is evaluated at these nodes. The quadrature method used in Uncertainpy is Leja quadrature, with Smolyak sparse grids to reduce the number of nodes required (Narayan and Jakeman, 2014; Smolyak, 1963). Pseudo-spectral projection methods are only used in Uncertainpy when requested by the user.

Several of the statistical metrics of interest can be obtained directly from the polynomial chaos expansion *Û*. The mean is simply

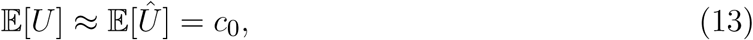

and the variance is

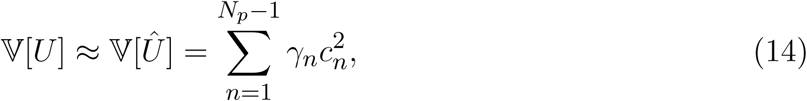

where *γ*_*n*_ is a normalization factor defined as

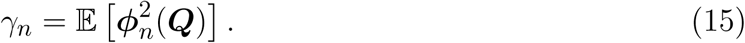

The first and total-order Sobol indices can also be calculated directly from the polynomial chaos expansion (Sudret, 2008; Crestaux et al., 2009). On the other hand, the percentiles and prediction interval must be estimated using *Û* as a surrogate model, and then perform the same procedure as for the (quasi-)Monte Carlo methods.

### 2.6 Dependency between uncertain parameters

One of the underlying assumptions when creating the polynomial chaos expansions is that the model parameters are independent. However, dependent parameters in neuroscience models are quite common (Achard and De Schutter, 2006). Fortunately, models containing dependent parameters can be analyzed with Uncertainpy by the aid of the Rosenblatt transformation from Chaospy (Rosenblatt, 1952; Feinberg and Langtangen, 2015). In brief, the idea is to create a reformulated model 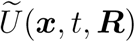 based on an independent parameter set ***R***, and then perform polynomial chaos expansions on the reformulated model. The Rosenblatt transformation is used to construct the reformulated model so it gives the same output (and statistics) as the original model, i.e.:

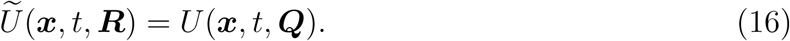

For more information on the use of the Rosenblatt transformation, see the Uncertainpy documentation^2^ or Feinberg and Langtangen (2015).

### 2.7 Feature-based analysis

When measuring the membrane potential of a neuron, the precise timing of action potentials often vary between recordings, even if the experimental conditions are maintained constant to the highest degree possible. This behaviour is typical for biological system. Since the experimental data displays such variation, it is often meaningless (or even misguiding) to base the success of a computational model on a direct point-to-point comparison between the experimental data and model output (Druckmann et al., 2007; Van Geit et al., 2008). A common modeling practice is therefore to rather have the model reproduce essential features of the experimentally observed dynamics, such as the action potential shape, or action potential firing rate (Druckmann et al., 2007). Such features are typically more robust between different experimental measurements, or between different model simulations, than the raw data or raw model output, at least if sensible features have been chosen.

Uncertainpy takes this aspect of neural modeling into account, and is constructed so it can extract a set of features relevant for various common model types in neuroscience from the raw model output. Examples include the action potential shape in single neuron models, or the mean interspike interval in neural network models. This means Uncertainpy performs an uncertainty quantification and sensitivity analysis not only on the raw model output, but also on a set of relevant features selected by the user. A list of the implemented features is given in Section 3.4, and the value of feature-based analysis is illustrated in two of the case studies (Section 4.3 and Section 4.4).

## 3. USER GUIDE FOR UNCERTAINPY

Uncertainpy is a Python toolbox, tailored to make uncertainty quantification and sensitivity analysis easily accessible to the computational neuroscience community. The toolbox is based on Python, since Python is a high level, open-source language in extensive and increasing use within the scientific community (Oliphant, 2007; Einevoll, 2009; Muller et al., 2015). Uncertainpy utilizes the Python-package Chaospy (Feinberg and Langtangen, 2015) to perform the uncertainty calculations. Uncertainpy currently only works with Python 2, due to limitations in some of the Python packages utilized. In this section we present a guide to how to use Uncertainpy. We do not present an exhaustive overview, but refer to the online description of Uncertainpy for a complete documentation. A complete case study with code is shown in Section 4.1.

Uncertainpy is easily installed by following the instructions in Section 3.8. After installation, we get access to Uncertainpy by simply importing it:

`import uncertainpy as un`

Performing an uncertainty quantification and sensitivity analysis with Uncertainpy includes three main components:

1. The **model** we want to examine.
2. The **parameters** of the model.
3. Specifications of **features** in the model output.

The model and parameters are required components, while the feature specifications are optional. The three (or two) components are brought together in the `UncertaintyQuantification` class. This class performs the uncertainty calculations and is the main class the user interacts with. In this section we explain how to use `UncertaintyQuantification` with the above components, and introduce a few additional utility classes.

### 3.1 The uncertainty quantification class

The `UncertaintyQuantification` class is used to define the problem, perform the uncertainty quantification and sensitivity analysis, and save and visualize the results. Among others, `UncertaintyQuantification` takes the arguments:

~~~
UQ = un. UncertaintyQuantification(
         model= Model (…),                *# Required*
         parameters= Parameters (…),      *# Required*
         features= Features (…)           *# Optional*
)
~~~

The arguments are given as instances of their corresponding Uncertainpy classes (`Model`, `Parameters`, and `Features`). We go through how to use each of these classes in the next three sections (Sections 3.2 to 3.4).

After the problem is set up, an uncertainty quantification and sensitivity analysis can be performed by using the `UncertaintyQuantification.quantify` method. Among others, `quantify` takes the optional arguments:

~~~
data = UQ. quantify(
    method=” pc"|” mc",
    pc_method=” collocation"|” spectral",
    rosenblatt= False| True
~~~

The `method` argument allows the user to choose whether Uncertainpy should use polynomial chaos expansions (“pc”) or quasi-Monte Carlo (“mc”) methods to calculate the relevant statistical metrics. If polynomial chaos expansions are chosen, `pc_method` further specifies whether point collocation (“collocation”) or spectral projection (“spectral”) methods are used to calculate the expansion coefficients. Finally, `rosenblatt` (False or True) determines if the Rosenblatt transformation should be used. If nothing is specified, Uncertainpy by default uses polynomial chaos expansions based on point collocation without the Rosenblatt transformation.

The results from the uncertainty quantification are returned in `data`, as a `Data` object (see Section 3.6). The results are also automatically saved in a folder named `data`, and figures are automatically plotted and saved in a folder named `figures`, both in the current directory. The returned `data` object is therefore not necessarily needed.

Polynomial chaos expansions are recommended as long as the number of uncertain parameters is small (typically > 20), as polynomial chaos expansions in these cases are much faster than quasi-Monte Carlo methods. Additionally, sensitivity analysis is not yet available for studies based on the quasi-Monte Carlo method. Which of the polynomial chaos expansion methods to choose is problem dependent, but in general the pseudo-spectral method is faster than point collocation, but has lower stability. We therefore generally recommend the point collocation method.

We note that there is no guarantee each set of sampled parameters produces a valid model or feature output. For example, a feature such as the spike width will not be defined in a model evaluation that produces no spikes. In such cases, Uncertainpy gives a warning which includes the number of runs that failed to return a valid output, and performs the uncertainty quantification and sensitivity analysis using the reduced set of valid runs. Point collocation (as well as the quasi-Monte Carlo method) are robust towards missing values as long as the number of results remaining is high enough (Eck et al., 2016), another reason the point collocation method is recommended. However, if a large fraction of the simulations fail, the user could consider redefining the problem (e.g., by using narrower parameter distributions).

### 3.2 Models

In order to perform the uncertainty quantification and sensitivity analysis of a model, Uncertainpy needs to set the parameters of the model, run the model using those parameters, and receive the model output. Uncertainpy has built-in support for NEURON and NEST models, found in the `NeuronModel` (Section 3.2.4) and `NestModel` (Section 3.2.5) classes respectively. It should be noted that while Uncertainpy is tailored towards neuroscience, it is not restricted to neuroscience models. Uncertainpy can be used on any model that meets the criteria in this section. Below, we first explain how to create custom models, before we explain how to use `NeuronModel` and `NestModel`.

#### 3.2.1 The model class

Generally, models are created through the `Model` class. `Model` takes the argument `run` and the optional arguments `postprocess`, `adaptive` and `labels`.

~~~
model = un. Model(run= example_model,
         postprocess= example_postprocess,
         adaptive= True,
         labels =[” xlabel", “ ylabel"])
~~~

The `run` argument must be a Python function that runs a simulation on a specific model for a given set of model parameters, and returns the simulation output. In this paper we call such a function for a model function. The `postprocess` argument is a Python function used to `postprocess` the model output if required. We go into details on the requirements of the postprocess and model functions below. The `adaptive` argument specifies whether the model uses adaptive time steps or not. For adaptive models, Uncertainpy automatically interpolates the output to a regular form (the same number of measurement points for each model evaluation, most commonly time points). Finally, `labels` allows the user to specify a list of labels to be used on the axes when plotting the results.

#### 3.2.2. Defining a model function

As explained above, the `run` argument is a Python function that runs a simulation on a specific model for a given set of model parameters, and returns the simulation output. An example outline of a model function is:

~~~
def example_model(parameter_1, parameter_2):
 *# An algorithm for the model, or a script that runs*
 *# an external model, using the given input parameters.*
 *# Returns the model output and model time*
 *# along with the optional info object.*
 return time, values, info
~~~

Such a model function has the following requirements:

1. **Input.** The model function takes a number of arguments which define the uncertain parameters of the model.
2. **Run the model.** The model must then be run using the parameters given as arguments.
3. **Output.** The model function must return at least two objects, the model time (or equivalent, if applicable) and model output. Additionally, any number of optional info objects can be returned. In Uncertainpy, we refer to the `time` object as time, the model output object as values, and the remaining objects as `info`.

a. **Time** (`time`). `time` can be interpreted as the *x*-axis of the model. It is used when interpolating (see below), and when certain features are calculated. We can return None if the model has no time associated with it.
b. **Model output** (`values`). The model output must either be regular, or it must be possible to interpolate or postprocess the output (see Section 3.2.3) to a regular form.
c. **Additional info** (`info`). Some of the methods provided by Uncertainpy, such as the later defined model postprocessing, feature preprocessing, and feature calculations, require additional information from the model (e.g., the time a neuron receives an external stimulus). We recommend to use a single dictionary as info object, with key-value pairs for the information, to make debugging easier. Uncertainpy always uses a single dictionary as the `info` object. Certain features require that specific keys are present in this dictionary.

The model itself does not need to be implemented in Python. Any simulator can be used, as long as we can control the model parameters and retrieve the simulation output via Python. We can as a shortcut pass a model function to the `model` argument in `UncertaintyQuantification`, instead of first having to create a `Model` instance.

#### 3.2.3 Defining a postprocess function

The `postprocess` function is used to postprocess the model output before it is used in the uncertainty quantification. Postprocessing does not change the model output sent to the feature calculations. This is useful if we need to transform the model output to a regular result for the uncertainty quantification, but still need to preserve the original model output to reliably detect the model features. Figure 2 illustrates how the objects returned by the model function are sent to both model `postprocess`, and feature `preprocess` (see Section 3.4).

**Figure 2.**
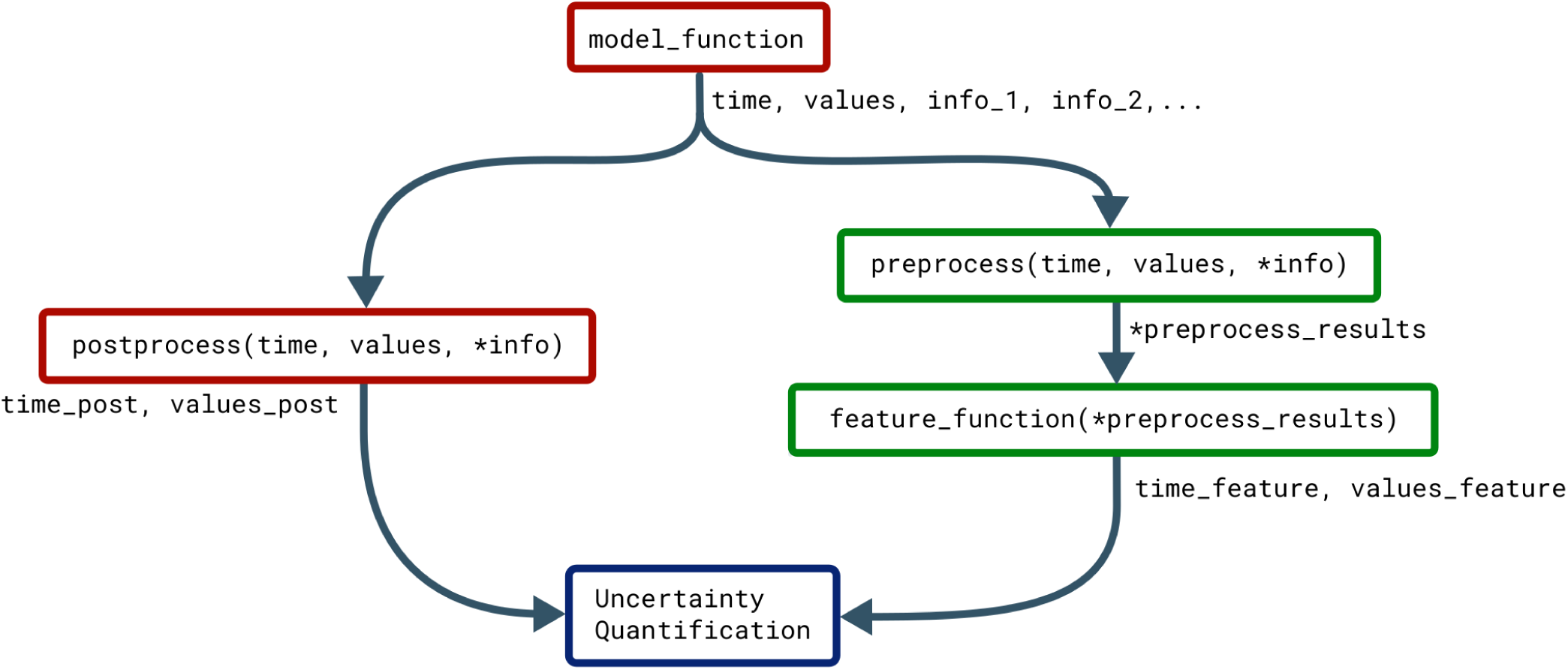
Classes that affect the objects returned by the model. The Uncertainpy methods that use, change, and perform calculations on the objects returned by the model function (time, values, and the optional info). Functions associated with the model are in red while functions associated with features are in green.

An example outline of the `postprocess` function is:

~~~
def example_postprocess(time, values, info):
*# Postprocess the result to a regular form using time,*
*# values, and info returned by the model function.*
*# Return the postprocessed model output and time.*
return time_postprocessed, values_postprocessed
~~~

The only time postprocessing is required for Uncertainpy to work, is when the model produces output that can not be interpolated to a regular form by Uncertainpy. Postprocessing is for example required for network models that give output in the form of spike trains, i.e. time values indicating when a given neuron fires. It should be noted that postprocessing of spike trains is already implemented in Uncertainpy (see Section 3.2.5). For most purposes user defined postprocessing will not be necessary.

The requirements for the `postprocess` function are:

A. **Input.** The `postprocess` function must take the objects returned by the model function as input arguments.
B. **Postprocessing.** The model time (`time`) and output (`values`) must be postprocessed to a regular form, or to a form that can be interpolated to a regular form by Uncertainpy. If additional information is needed from the model, it can be passed along in the `info` object.
C. **Output.** The `postprocess` function must return two objects:

a. **Model time** (`time_postprocessed`). The first object is the postprocessed time (or equivalent) of the model. We can return `None` if the model has no time. Note that the automatic interpolation of the postprocessed time can only be performed if a postprocessed time is returned (if an interpolation is required).
b. **Model output** (`values_postprocessed`). The second object is the postprocessed model output.

#### 3.2.4 NEURON model class

NEURON (Hines and Carnevale, 1997) is a widely used simulator for multi-compartmental neural models. Uncertainpy has support for NEURON models through the NeuronModel class, a subclass of Model. Among others, `NeuronModel` takes the arguments:

~~~
model = un. NeuronModel(path=” path/ to/ neuron_model",
            adaptive= True,
            stimulus_start =1000,                       *# ms*
            stimulus_end =1900)                         *# ms*
~~~

`path` is the path to the folder where the NEURON model is saved (the location of the mosinit.hoc file). `adaptive` indicates whether the NEURON model uses adaptive time steps. stimulus_start and stimulus_end denotes the start and end time of any stimulus given to the neuron. `NeuronModel` loads the NEURON model from mosinit.hoc, sets the parameters of the model, evaluates the model and returns the somatic membrane potential of the neuron. `NeuronModel` therefore does not require a model function. A case study of a NEURON model analysed with Uncertainpy is found in Section 3.2.4.

If changes are needed to the standard `NeuronModel`, such as measuring the voltage from other locations than the soma, the `Model` class with an appropriate model function should be used instead. Alternatively, `NeuronModel` can be subclassed and the existing methods customized as required. An example of the later is shown in `/uncertainpy/examples/bahl/`.

#### 3.2.5 NEST model class

NEST (Eppler et al., 2015) is a simulator for large networks of spiking neurons. NEST models are supported through the `NestModel` class, another subclass of `Model`:

~~~
model = un. NestModel(run= nest_model_function)
~~~

`NestModel` requires the model function to be specified through the `run` argument, unlike `NeuronModel`. The NEST model function has the same requirements as a regular model function, except it is restricted to return only two objects: the final simulation time (denoted `simulation_end`), and a list of spike times for each neuron in the network, which we refer to as spike trains (denoted `spiketrains`).

A spike train returned by a NEST model is a set of irregularly spaced time points where a neuron fired a spike. NEST models therefore require postprocessing to make the model output regular. Such a postprocessing is provided by the implemented `NestModel.postprocess` method, which converts a spike train to a list of zeros (no spike) and ones (a spike) for each time step in the simulation. For example, if a NEST simulation returns the spike train [0, 2, 3.5], it means the neuron fired three spikes occurring at *t* = 0, 2, and 3.5 ms. If the simulation have a time resolution of 0.5 ms and ends after 4 ms, `NestModel.postprocess` returns the postprocessed spike train `[1, 0, 0, 0, 1, 0, 0, 1, 0]`, and the postprocessed time array `[0, 0.5, 1, 1.5, 2, 2.5, 3, 3.5, 4]`. The final uncertainty quantification of a NEST network therefore predicts the probability for a spike to occur at any specific time point in the simulation. An Uncertainpy-based analysis of a NEST model is found in the case study in Section 4.4.

### 3.3. Parameters of the model

The parameters of a model are defined by two properties, they must have (i) a name and either a fixed value or a distribution. It is important that the name of the parameter is the same as the name given as the input argument in the model function. A parameter is considered uncertain if it is given a probability distribution, and the distributions are given as Chaospy distributions. 64 different univariate distributions are defined in Chaospy, and Chaospy has support for easy creation of multivariate distributions. For a list of available distributions and detailed instructions on how to create probability distributions with Chaospy, see Section 3.3 in Feinberg and Langtangen (2015).

The parameters are defined by the `Parameters` class. `Parameters` takes the argument `parameters`, which is a dictionary with the above information. The names of the parameters are the keys, and the fixed values or distributions of the parameters are the values. As an example, if we have two parameters, where the first is named `name_1` and has a uniform probability distributions in the interval [8, 16], and the second is named `name_2` and has a fixed value 42, the list becomes:

~~~
import chaospy as cp
parameters = {” name_1 “: cp. Uniform (8, 16), “ name_2 “: 42}
~~~

And `Parameters` is initialized:

~~~
parameters = un. Parameters(parameters = parameters)
~~~

We can as a shortcut pass the above dictionary to the `parameters` argument in `UncertaintyQuantification`, instead of first having to create a `Parameters` instance.

If the parameters do not have separate univariate probability distributions, but a joint multivariate probability distribution, the multivariate distribution can be set by giving `Parameters` the optional argument `distribution`:

~~~
multivariate = cp.J(cp. Uniform (8, 16), cp. Uniform (40, 44))
parameters = un.Parameters(parameters= parameters,
                          distribution = multivariate))
~~~

### 3.4 Features

As discussed in Section 2.7, it is often more meaningful to examine the uncertainty in salient features of the model output, than to base the analysis directly on a point-to-point comparison of the raw output data (e.g. a voltage trace). Upon user request, Uncertainpy can identify and extract features of the model output. If we give the `features` argument to `UncertaintyQuantification`, Uncertainpy will perform uncertainty quantification and sensitivity analysis of the given features, in addition to the analysis of the raw output data (if desired).

Three sets of features comes predefined with Uncertainpy, `SpikingFeatures`, `EfelFeatures`, and `NetworkFeatures`. Each of the feature classes contains a set of features tailored towards one specific type of models. We first explain how to create custom features, before explaining how to use the built-in features.

Features are defined through the `Features` class:

~~~
list_of_feature_functions = [ example_feature]

features = un.Features(new_features= list_of_feature_functions,
                      features_to_run =[” example_feature"],
                      preprocess= example_preprocess,
                      adaptive =[” example_feature"])
~~~

`new_features` is a list of Python functions that each calculates a specific feature, whereas `features_to_run tells` which of the features to perform uncertainty quantification of. If nothing is specified, the uncertainty quantification is by default performed on all features (`features_to_run="all"`). `preprocess` is a Python function that performs common calculations for all features. adaptive is a list of features that have adaptive time steps. As with models, Uncertainpy automatically interpolates the output of adaptive features to a regular form. Below we first go into details on the requirements of a feature function, and then the requirements of a `preprocess` function.

#### 3.4.1 Feature functions

A specific feature is given as a Python function. The outline of such a feature function is:

~~~
def example_feature(time, values, info):
  *# Calculate the feature using time, values and info.*
  *# Return the feature times and values.*
return time_feature, values_feature
~~~

Feature functions have the following requirements:

1. **Input.** The feature function takes the objects returned by the model function as input, except in the case when a `preprocess` function is used (see below). In that case, the feature function instead takes the objects returned by the `preprocess` function as input. `preprocess` is normally not used.
2. **Feature calculation.** The feature function calculates the value of a feature from the data given in time, values and optional `info` objects. As previously mentioned, in all built-in features in Uncertainpy, `info` is a dictionary containing required information as key-value pairs.
3. **Output.** The feature function must return two objects:

a. **Feature time** (`time_feature`). The time (or equivalent) of the feature. We can return

None instead for features where it is not relevant.

a. **Feature values** (`values_feature`). The result of the feature calculation. As for the model output, the feature results must be regular, or able to be interpolated. If there are no feature results for a specific model evaluation (e.g., if the feature was spike width and there was no spike), the feature function can return `None`. The specific feature evaluation is then discarded in the uncertainty calculations.

As with models, we can as a shortcut give a list of feature functions as the `feature` argument in `UncertaintyQuantification`, instead of first having to create a `Features` instance.

#### 3.4.2 Feature preprocessing

Some of the calculations needed to quantify features may overlap between different features. One example is finding the spike times from a voltage trace. The `preprocess` function is used to avoid having to perform the same calculations several times. An example outline of a `preprocess` function is:

~~~
def preprocess(time, values, info):
   *# Perform all common feature calculations using time,*
   *# values, and info returned by the model function.*

   *# Return the preprocessed model output and info.*
   return time_preprocessed, values_preprocessed, info
~~~

The requirements for a `preprocess` function are:

1. **Input.** A `preprocess` function takes the objects returned by the model function as input.
2. **Preprocesssing.** The model output `time, values`, and additional `info` objects are used to perform all `preprocess` calculations.
3. **Output.** The `preprocess` function can return any number of objects as output. The returned `preprocess` objects are used as input arguments to the feature functions, so the two must be compatible.

Figure 2 illustrates how the objects returned by the model function are passed to preprocess, and the returned `preprocess` objects are used as input arguments in all feature functions. The preprocessing makes it so feature functions have different required input arguments depending on the feature class they are added to. As mentioned earlier, Uncertainpy comes with three built-in feature classes. These classes all take the `new_features` argument, so custom features can be added to each set of features. These feature classes all perform a preprocessing, and therefore have different requirements for the input arguments of new feature functions. Additionally, certain features require specific keys to be present in the `info` dictionary. Each class has a `reference_feature` method that states the requirements for feature functions of that class in its docstring.

#### 3.4.3 Spiking features

Here we introduce the `SpikingFeatures` class, which contains a set of features relevant for models of single neurons that receive an external stimulus and responds by eliciting a series of action potentials, also called spikes. Many of these features require the start time and end time of the stimulus, which must be returned as `info["stimulus_start"]` and `info["stimulus_start"]` in the model function. `info` is then used as an additional input argument in the calculation of each feature. A set of spiking features is created by:

~~~
features = SpikingFeatures ()
~~~

`SpikingFeatures` implements a `preprocess` method, which locates spikes in the model output. This `preprocess` can be customized, see the documentation on `SpikingFeatures`.

The features included in `SpikingFeatures` are briefly defined below. This set of features was taken from the previous work of Druckmann et al. (2007), with the addition of the number of action potentials during the stimulus period. We refer to the original publication for more detailed definitions.

1. `nr_spikes` – Number of action potentials (during stimulus period).
2. `spike_rate` – Action-potential firing rate (number of action potentials divided by stimulus duration).
3. `time_before_first_spike` – Time from stimulus onset to first elicited action potential.
4. `accommodation_index` – Accommodation index (normalized average difference in length of two consecutive interspike intervals).
5. `average_AP_overshoot` – Average action-potential peak voltage.
6. `average_AHP_depth` – Average afterhyperpolarization depth (average minimum voltage between action potentials).
7. `average_AP_width` – Average action potential width taken at midpoint between the onset and peak of the action potential.

The user may want to add custom features to the set of features in `SpikingFeatures`. The `SpikingFeatures`.preprocess method changes the input given to the feature functions, and as such each spiking feature function has the following input arguments:

1. The `time` array returned by the model simulation.
2. An `Spikes` object (spikes) which contain the spikes found in the model output.
3. An `info` dictionary with `info["stimulus_start"]` and `info["stimulus_end"]` set.

The `Spikes` object is the preprocessed version of the model output, used as a container for Spike objects. In turn, each Spike object contain information of a single spike. This information includes a brief voltage trace represented by a `time` and a voltage (V) array that only includes the selected spike. The information in `Spikes` is used to calculate each feature. As an example, let us assume we want to create a feature that is the time at which the first spike in the voltage trace ends. Such a feature can be defined as follows:

~~~
def first_spike_end_time(time, spikes, info):
  *# Calculate the feature from the spikes object*
  spike = spikes [0]                    *# Get the first spike*
  values_feature = spike.t[-1]          *# The last time point in the spike*

  return None, values_feature
~~~

This feature may now be used as a feature function in the list given to the `new_features` argument.

From the set of both built-in and user defined features, we may select subsets of features that we want to use in the analysis of a model. Let us say we are interested in how the model performs in terms of the three features: `nr_spikes, average_AHP_depth` and `first_spike_end_time`. A spiking features object that calculates these features is created by:

~~~
features_to_run = [” nr_spikes",
                   ” average_AHP_depth",
                   ” first_spike_end_time"]
features = un. SpikingFeatures(new_features=[ first_spike_end_time],
                                                     features_to_run= features_to_run)
~~~

#### 3.4.4 Efel features

A more extensive set of features for single neuron voltage traces is found in the Electrophys Feature Extraction Library (eFEL) (Blue Brain Project, 2015). A set of eFEL spiking features is created by:

`features = EfelFeatures ()`

Uncertainpy has all features in the eFEL library in the `EfelFeatures` class. eFEL currently contains 153 different features. Due to the high number of features, we do not list them here, but refer to the eFEL documentation^3^ for detailed definitions, or the Uncertainpy documentation for a list of the features. `EfelFeatures` is used in the same way as the `SpikingFeatures`.

#### 3.4.5 Network features

The last set of features implemented in Uncertainpy is found in the `NetworkFeatures` class:

~~~
features = NetworkFeatures ()
~~~

This class contains a set of features relevant for the output of neural network models and are calculated using the Elephant software (NeuralEnsemble, 2017). The implemented features are:

1. `mean_firing_rate` – Mean firing rate (averaged over all recorded neurons).
2. `instantaneous_rate` – Instantaneous firing rate (averaged over all recorded neurons within a small time window).
3. `mean_isi` – Mean interspike interval (averaged over all recorded neurons).
4. `cv` – Coefficient of variation of the interspike interval (for a single recorded neuron).
5. `mean_cv` – Mean coefficient of variation of the interspike interval (averaged over all recorded neurons).
6. `lv` – Local variation (variability of interspike intervals for a single recorded neuron).
7. `mean_lv` – Mean local variation (variability of interspike intervals averaged over all recorded neurons).
8. `fanofactor` – Fanofactor (variability of spike trains).
9. `victor_purpura_dist` – Victor Purpura distance (spike train dissimilarity between two recorded neurons).
10. van_rossum_dist – Van Rossum distance (spike train dissimilarity between two recorded neurons).
11. `binned_isi` – Histogram of the interspike intervals (for all recorded neurons).
12. `corrcoef` – Pairwise Pearson’s correlation coefficients (between the binned spike trains of two recorded neurons).
13. `covariance` – Covariance (between the binned spike trains of two recorded neurons).

A few of these network features can be customized, see the documentation on `NetworkFeatures` for a further explanation.

The use of `NetworkFeatures` in Uncertainpy follows the same logic as the use of the other feature classes, and custom features can easily be included. As with `SpikingFeatures`, `NetworkFeatures` implements a `preprocess` method. This `preprocess` returns the following objects:

1. End time of the simulation (`end_time`).
2. A list of NEO (Garcia et al., 2014) spike trains (`spiketrains`).

Each feature function therefore require the same objects as input arguments. Note that the `info` object is not used.

### 3.5 Uncertainty calculations in Uncertainpy

In this section we describe how Uncertainpy performs the uncertainty calculations, as well as which options the user have to customize the calculations. Moreover, a detailed insight into this is not required to use Uncertainpy, as in most cases, the default settings works fine. In addition to the customization options shown below, Uncertainpy has support for implementing entirely custom uncertainty quantification and sensitivity analysis methods. This is only recommended for expert users, as knowledge of both Uncertainpy and uncertainty quantification is needed. We do not go into details here, but refer to the Uncertainpy documentation for more information.

#### 3.5.1 Quasi-Monte Carlo method

To use the quasi-Monte Carlo method, we call `quantify` with `method="mc"`, and the optional argument `nr_mc_samples`:

~~~
data = UQ. quantify(
    method=” mc",
    nr_mc_samples =10**3,
)
~~~

By default, the quasi-Monte Carlo method quasi-randomly draws 1000 parameter samples from the joint multivariate probability distribution of the parameters *ρ***_*Q*_** using Hammersley sampling (Hammersley, 1960). As the name indicates, the number of samples is specified by the `nr_mc_samples` argument. The model is evaluated for each of these parameter samples, and features are calculated for each model evaluation (when applicable). To speed up the calculations, Uncertainpy uses the *multiprocess* Python package (McKerns et al., 2012) to perform this step in parallel. When model and feature calculations are done, Uncertainpy calculates the mean, variance, and 5-th and 95-th percentile (which gives the 90% prediction interval) for the model output as well as for each feature.

#### 3.5.2 Polynomial chaos expansions

To use polynomial chaos expansions we use `quantify` with the argument `method="pc"`, which takes a set of optional arguments (the specified values are default):

~~~
data = UQ. quantify(method=” pc",
    pc_method=” collocation",
    rosenblatt= False,
    polynomial_order =3,
    nr_collocation_nodes= None,
    quadrature_order= None,
    nr_pc_mc_samples =10**4,
)
~~~

As previously mentioned, Uncertainpy allows the user to select between point collocation (`pc_method="collocation”`) and pseudo-spectral projections (`pc_method="spectral”`). The goal is to create separate polynomial chaos expansions *Û* for the model and each feature. In both methods, Uncertainpy creates the orthogonal polynomial ***ϕ****n* using the three-term recurrence relation and *ρ***_*Q*_**. Uncertainpy uses a third order polynomial expansion, which can be changed with the `polynomial_order` argument. The polynomial ***ϕ****n* is shared between the model and all features, since they have the same uncertain input parameters, and therefore the same *ρ***_*Q*_**. Only the polynomial coefficients *c*_*n*_ differ between the model and each feature.

The two polynomial chaos methods differ in terms of how they calculate *c*_*n*_. For point collocation Uncertainpy uses 2(*N*_*p*_ + 1) collocation nodes, as recommended by Hosder et al. (2007), where *N*_*p*_ is the number of polynomial chaos expansion factors. The number of collocation nodes can be customized with `nr_collocation_nodes`, but the new number of nodes must be chosen carefully. The collocation nodes are sampled from *ρ***_*Q*_** using Hammersley sampling (Hammersley, 1960). The model and features are calculated for each of the collocation nodes. As with the quasi-Monte Carlo method, this step is performed in parallel. The polynomial coefficients *c*_*n*_ are calculated using the model and feature results, and Tikhonov regularization (Rifkin and Lippert, 2007).

For the pseudo-spectral projection, Uncertainpy chooses nodes and weights using a quadrature scheme, instead of choosing nodes from *ρ***_*Q*_**. The quadrature scheme used is Leja quadrature with a Smolyak sparse grid (Narayan and Jakeman, 2014; Smolyak, 1963). The Leja quadrature is of order two greater than the polynomial order, but can be changed with `quadrature_order`. The model and features are calculated for each of the quadrature nodes. As before, this step is performed in parallel. The polynomial coefficients *c*_*n*_ are then calculated from the quadrature nodes, weights, and model and feature results.

When Uncertainpy has derived *Û* for the model and features, it uses *Û* to compute the mean, variance, and the first and total-order Sobol indices. The first and total-order Sobol indices are also summed and normalized. Finally, Uncertainpy uses *Û* as a surrogate model, and performs a quasi-Monte Carlo method with Hammersley sampling and `nr_pc_mc_samples=10**4` samples to find the 5-th and 95-th percentiles.

If the model parameters have a dependent joint multivariate distribution, the Rosenblatt transformation must be used by setting `rosenblatt=True`. To perform the transformation Uncertainpy chooses *ρ***_*R*_** to be a multivariate independent normal distribution, which is used instead of *ρ***_*Q*_** to perform the polynomial chaos expansions. Both the point collocation method and the pseudo-spectral method are performed as described above. The only difference is that we use *ρ***_*R*_** instead of *ρ***_*Q*_**, and use the Rosenblatt transformation to transform the selected nodes from ***R*** to ***Q***, before they are used in the model evaluation.

### 3.6 Data format

All results from the uncertainty quantification and sensitivity analysis in Uncertainpy is returned as a `Data` object, as well as being stored in `UncertaintyQuantification.data`. The `Data` class works similarly to a Python dictionary. The name of the model or feature is the key, while the values are `DataFeature` objects that stores each statistical metric in Table 1 as attributes. Results can be saved and loaded through `Data.save` and `Data.load`, and are saved as `hdf5` files (Collette, 2013).

**Table 1.** Calculated values and statistical metrics, for the model and each feature stored in the `Data` class.

| Calculated statistical metric | Symbol | Variable |
| --- | --- | --- |
| Model and feature evaluations | $U$ | `evaluations` |
| Model and feature times | $t$ | `time` |
| Mean | $\mathbb{E}$ | `mean` |
| Variance | $\mathbb{V}$ | `variance` |
| 5-th percentile | $P_5$ | `percentile_5` |
| 95-th percentile | $P_{95}$ | `percentile_95` |
| First-order Sobol indices | $S$ | `sobol_first` |
| Total-order Sobol indices | $S_T$ | `sobol_total` |
| Normalized sum of first-order Sobol indices | $\hat{S}$ | `sobol_first_sum` |
| Normalized sum of total-order Sobol indices | $\hat{S}_T$ | `sobol_total_sum` |

An example: if we have performed uncertainty quantification of a spiking neuron model with the number of spikes as one of the features, we can load the results and get the variance of the number of spikes by typing:

~~~
data = un. Data()
data. load(“ filename”)
variance = data[” nr_spikes"]. variance
~~~

### 3.7 Visualization

Uncertainpy plots the results for all zero and one-dimensional statistical metrics, and some two-dimensional statistical metrics. These visualizations are intended as quick way to get an overview of the results, and not to create publication-ready plots. Custom plots of the data can easily be created by retrieving the results from the `Data` class.

### 3.8 Technical aspects

Uncertainpy is open-source and found at https://github.com/simetenn/uncertainpy. Uncertainpy can easily be installed using `pip`:

~~~
pip install uncertainpy
~~~

or from source by cloning the Github repository:

~~~
$ git clone https://github.com/simetenn/uncertainpy
$ cd uncertainpy
$ sudo python setup. py install
~~~

Uncertainpy comes with an extensive test suite that can be run with the `test.py` script. For information on how to use `test.py`, run:

~~~
$ python test. py -- help
~~~

## 4

EXAMPLE APPLICATIONS

In the current section, we demonstrate how to use Uncertainpy by applying it to four different case studies: (i) a simple model for the temperature of a cooling coffee cup implemented in Python, (ii) the original Hodgkin-Huxley model implemented in Python, a multi-compartment model of a thalamic interneuron implemented in NEURON, and a sparsely connected recurrent network model implemented in NEST. All four case studies are available in `/uncertainpy/examples/`, which generates all results shown in this paper. All case studies can be run on a regular workstation computer. Uncertainpy does not create publication-ready figures, so custom plots have been created for the case studies below. The code for creating all figures in this paper is found in a Jupyter Notebook in `/uncertainpy/examples/article_plots/`. The version of Uncertainpy used to create these results is commit `2503fec`.

### 4.1 Cooling coffee cup

To give a simple, first demonstration of Uncertainpy, we perform an uncertainty quantification and sensitivity analysis of a hot cup of coffee that follows Newton’s law of cooling. We start with a model that has independent uncertain parameters, before we expand the model to have dependent parameters to show the use of the Rosenblatt transformation.

#### 4.1.1 Cooling coffee cup with independent parameters

The temperature *T* of the cooling coffee cup is given by:

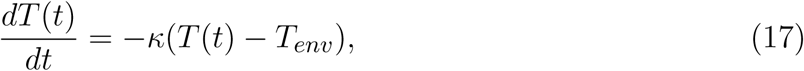

where *T*_*env*_ is the temperature of the environment in units of °C. *κ* is a cooling constant in units of 1/min that is characteristic of the system and regulates how fast the coffee cup radiates heat to the environment. We set the initial temperature to a fixed value, *T*_0_ = 95°C, and assume that *κ* and *T*_*env*_ are uncertain parameters characterized by the uniform probability distributions:

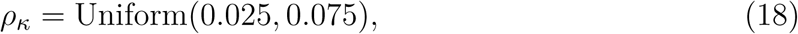

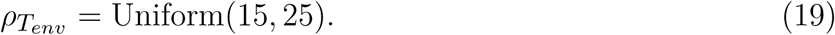

The following code is available in `/uncertainpy/examples/coffee_cup/`. We start by importing the packages required to perform the uncertainty quantification:

~~~
import uncertainpy as un
import chaospy as cp                    *# To create distributions*
import numpy as np                      *# For the time array*
from scipy. integrate import odeint     *# To integrate our equation*
~~~

Next, we create the cooling coffee-cup model. To do this we define a Python function (coffee_cup) that takes the uncertain parameters kappa and T_env as input arguments, solves Equation (17) by integration using scipy.integrate.odeint over 200 minutes, and returns the resulting time and temperature arrays.

~~~
def coffee_cup(kappa, T_env):
  *# Initial temperature and time array*
  time = np. linspace (0, 200, 150)      *# Minutes*
  T_0 = 95                               *# Celsius*
*# The equation describing the model*
def f(T, time, kappa, T_env):
    return - kappa *(T - T_env)
*# Solving the equation by integration.*
temperature = odeint(f, T_0, time, args =(kappa, T_env))[:, 0]

*# Return time and model output*
return time, temperature
~~~

We now use `coffee_cup` to create a `Model` object, and add labels:

~~~
model = un.Model(run= coffee_cup,
         labels =[” Time (min)", “ Temperature (C)"])
~~~

As previously mentioned, it is possible to use `coffee_cup` directly as the `model` argument in the `UncertaintyQuantification` class, however we would then be unable to specify the labels.

In the next step, we use Chaospy to assign distributions to the uncertain parameters *κ* and *T*_*env*_, and use these distributions to create a `Parameters` object:

~~~
*# Create the distributions*
kappa_dist = cp. Uniform (0.025, 0.075)
T_env_dist = cp. Uniform (15, 25)

*# Define the parameters dictionary*
parameters = {” kappa": kappa_dist, “ T_env": T_env_dist}

*# and use it to create the parameters*
parameters = un. Parameters(parameters)
~~~

We can now set up the `UncertaintyQuantification`:

~~~
UQ = un.UncertaintyQuantification(model= model,
                           parameters= parameters)
~~~

With that, we are ready to calculate the uncertainty and sensitivity of the model. We use polynomial chaos expansions with point collocation, the default options of `quantify:`

~~~
data = UQ. quantify ()
~~~

`quantify` calculates all statistical metrics discussed in Section 2.2 and Section 2.3, but here we only show the mean, variance, and 90% prediction interval (Figure 3A), and the first-order Sobol indices (Figure 3B). As the mean (blue line) in Figure 3A shows, the cooling gives rise to an exponential decay in the temperature, towards the temperature of the environment *T*_env_. From the sensitivity analysis (Figure 3B) we see that *T* is most sensitive to *κ* early in the simulation, and to *T*_env_ towards the end of the simulation. This is as expected, since *κ* determines the rate of the cooling, while *T*_env_ determines the final temperature. After about 150 minutes, the cooling is essentially completed, and the uncertainty in *T* exclusively reflects the uncertainty of *T*_env_.

**Figure 3.**
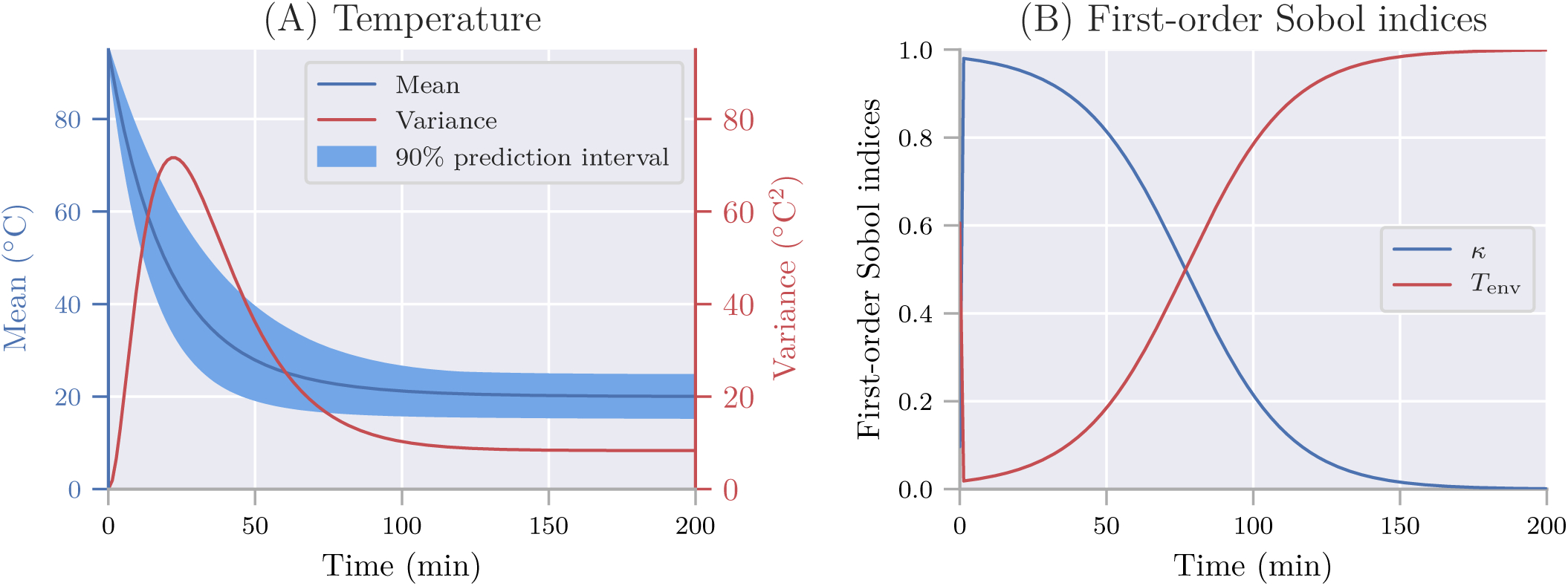
The uncertainty quantification and sensitivity analysis for the cooling coffee-cup model. (A) The mean, variance and 90% prediction interval of the temperature of the cooling coffee cup. (B) First-order Sobol indices of the cooling coffee-cup model.

#### 4.1.2 Cooling coffee cup with statistically dependent parameters

Uncertainpy can also perform uncertainty quantification and sensitivity analysis using polynomial chaos expansions on models with statistically dependent parameters. Here we use the cooling coffee-cup model to construct such an example. Let us parameterize the coffee cup differently:

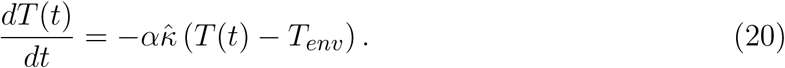

In order for the model to describe the same cooling process as before, the new variables *α* and 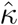 should be dependent, so 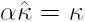 We can achieve this by demanding that 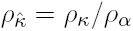 and otherwise define the problem following the same procedure as in the original case study. Since we now have dependent parameters, Uncertainpy returns an error message unless we tell it to use the Rosenblatt transformation to solve the problem:

~~~
data = UQ. quantify(rosenblatt= True)
~~~

In this case, the distribution we assign to *α* does not matter for the end result, as the distribution for 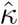 will be scaled accordingly. Using the Rosenblatt transformation, an uncertainty quantification and sensitivity analysis of the dependent coffee-cup model therefore returns the same results as seen in Figure 3, where the role of the original *κ* is taken over by 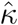 while the sensitivity to the additional parameter *α* becomes strictly zero (we do not show the results here, but see example in `/uncertainpy/examples/coffee_cup_dependent/`).

### 4.2 Hodgkin-Huxley model

From here on, we focus on case studies more relevant to neuroscience, starting with the original Hodgkin-Huxley model (Hodgkin and Huxley, 1952). An uncertainty analysis of this model has been performed previously (Torres Valderrama et al., 2015), and here we repeat a part of that study using Uncertainpy.

The here used version of the Hodgkin-Huxley model has 11 parameters with the numerical values listed in Table 2. As in the previous study, we assume each of these parameters have a uniform distribution in the range *±* 10% around their original value. We use uncertainty quantification and sensitivity analysis to explore how this parameter uncertainty affect the model output, i.e., the action potential response of the neural membrane potential *V*_*m*_ to an external current injection.

**Table 2.** Parameters in the original Hodgkin-Huxley model.

| Parameter | Value | Unit | Meaning |
| --- | --- | --- | --- |
| $V_0$ | -10 | mV | Initial voltage |
| $C_m$ | 1 | $\mu\text{F}/\text{cm}^2$ | Membrane capacitance |
| $\bar{g}_{\text{Na}}$ | 120 | $\text{mS}/\text{cm}^2$ | Sodium (Na) conductance |
| $\bar{g}_{\text{K}}$ | 36 | $\text{mS}/\text{cm}^2$ | Potassium (K) conductance |
| $\bar{g}_{\text{I}}$ | 0.3 | $\text{mS}/\text{cm}^2$ | Leak current conductance |
| $E_{\text{Na}}$ | 112 | mV | Sodium equilibrium potential |
| $E_{\text{K}}$ | -12 | mV | Potassium equilibrium potential |
| $E_{\text{I}}$ | 10.613 | mV | Leak current equilibrium potential |
| $n_0$ | 0.0011 | | Initial potassium activation gating variable |
| $m_0$ | 0.0003 | | Initial sodium activation gating variable |
| $h_0$ | 0.9998 | | Initial sodium inactivation gating variable |

As in the cooling coffee-cup example, we implement the Hodgkin-Huxley model as a Python function, and use polynomial chaos expansions with point collocation to calculate the uncertainty and sensitivity of the model (the code for this case study is found in `/uncertainpy/examples/valderrama/`).

The uncertainty quantification of the Hodgkin-Huxley model is shown in Figure 4A, and the sensitivity analysis in Figure 4B. Although the action potential is robust (within the selected parameter ranges), the onset and amplitude of the action potential varied. The variance in *V*_*m*_ is largest during the upstroke and peak of the action potential (Figure 4A), which occurs in the time interval between *t* = 8 and 9 ms.

**Figure 4.**
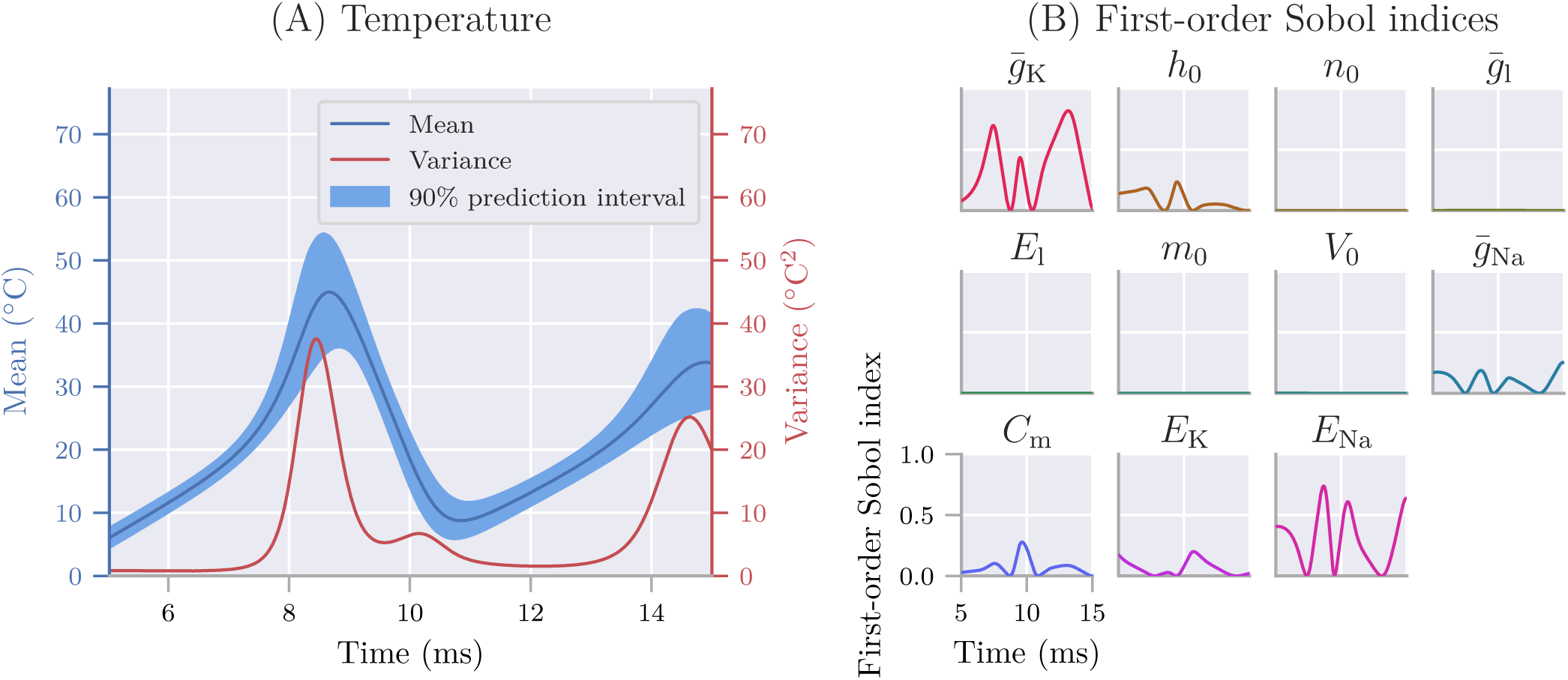
The uncertainty quantification and sensitivity analysis for the Hodgkin-Huxley model. The model was exposed to a continuous external stimulus of 140 *µ*A/cm2 starting at *t* = 0, and we examined the membrane potential in the time window between *t* = 5 and 15 ms. (A) Mean, variance and 90% prediction interval for the membrane potential of the Hodgkin-Huxley model. (B) First-order Sobol indices of the uncertain parameters in the Hodgkin-Huxley model.

The sensitivity analysis reveals that the variance in *V*_*m*_ mainly is due to the uncertainty in two parameters: the maximum conductance of the K^+^ channel, 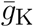 and the Na^+^ reversal potential, *E*_Na_ (Figure 4B). We see that 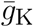 is most important during the upstroke of the action potential, indicating the K^+^ channel plays an important role for the timing of the action potential. *E*_Na_ is most important during the peak, reflecting the important role of *E*_Na_ in determining the action potential amplitude. The low sensitivity to the remaining parameters means that most of the variability of the Hodgkin-Huxley model would be maintained if these remaining parameters were kept fixed (at least for the given stimulus conditions). Due to some lacking implementation details in Torres Valderrama et al. (2015), our analysis is likely not an exact replica of the previous study, but the results obtained are quantitatively similar.

### 4.3 Multi-compartment model of a thalamic interneuron

In the next case study, we illustrate how Uncertainpy can be used on models implemented in NEURON (Hines and Carnevale, 1997). For this study, we select a previously published model of an interneuron in the dorsal lateral geniculate nucleus of the thalamus(Halnes et al., 2011). Since the model is implemented in NEURON, the original model can be used directly with Uncertainpy with the use of the `NeuronModel` class. The code for this case study is found in `/uncertainpy/examples/interneuron/`.

In the original modeling study, a set of 11 parameters were tuned manually through trial and error until the interneuron model obtained the desired response characteristics (Halnes et al., 2011). The final parameter set is listed in Table 3. To perform an uncertainty quantification and sensitivity analysis of this model, we assume each of these 11 parameters have a uniform uncertainty distribution in the interval *±*2.5% around their original value.

**Table 3.**
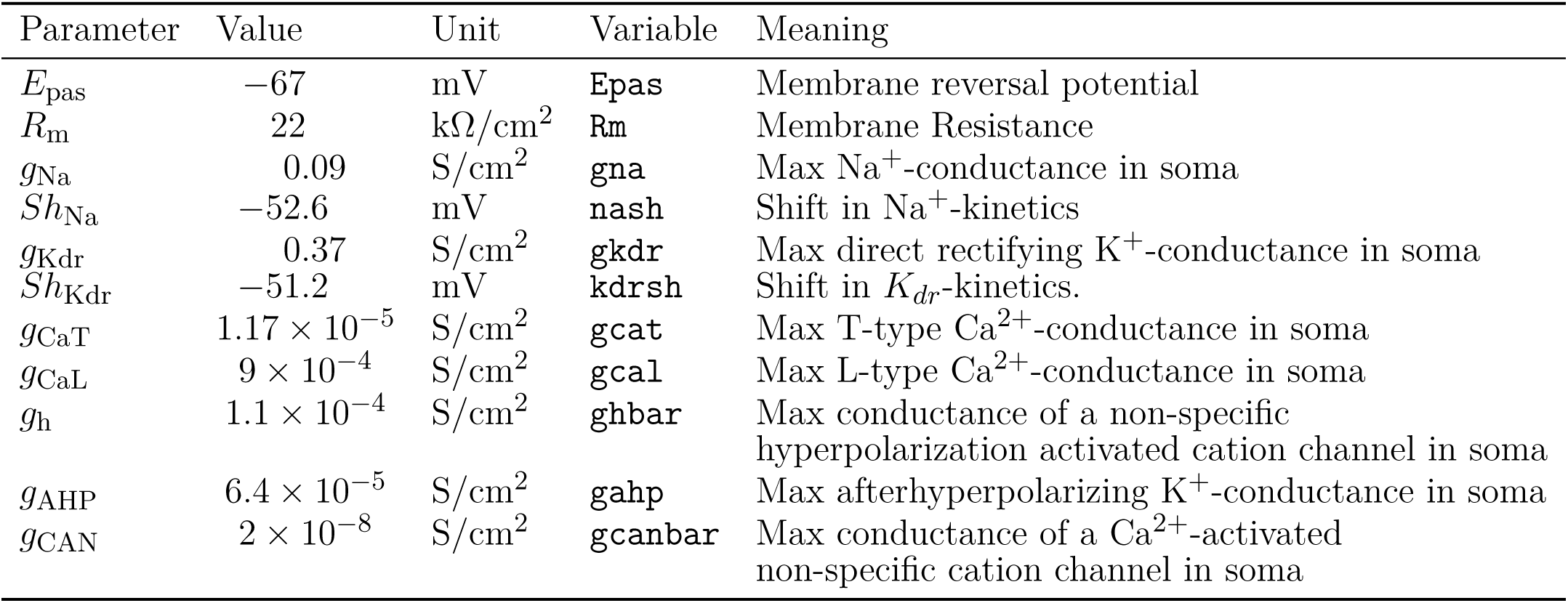
Uncertain parameters in the thalamic interneuron model.

As discussed in Section 2.7, a point-to-point comparison of voltage traces is often uninformative, and we therefore want to perform a feature-based analysis of the model. Since we examine a spiking neuron model, we choose the features in the `SpikingFeatures` class.

The simulation starts at *t* = 0 and we study the response of the interneuron to a somatic current injection between 1000 ms *< t <* 1900 ms. `SpikingFeatures` needs to know the start and end time of this stimulus to be able to calculate certain features. They are specified through the `stimulus_start` and `stimulus_end` arguments when initializing `NeuronModel`. Additionally, the interneuron model uses adaptive time steps, meaning we have to set `adaptive=True`. We also give the path to the folder where the neuron model is stored with `path="interneuron_modelDB/"`. As before, we use polynomial chaos expansion with point collocation to compute the statistical metrics for the model output and all features.

The uncertainty quantification of the membrane potential in the soma of the interneuron is seen in Figure 5A. The model is exposed to a somatic current injection and typically respond by eliciting one or several action potentials. To illustrate the variety of response characteristics hiding in the statistics in Figure 5A, four selected example simulations are shown in Figure 5B, all obtained by drawing the uncertain parameters from intervals 2.5% of their original values. The qualitative differences between the simulations in Figure 5B indicate that a feature-based analysis is more informative than a point-to-point comparison of the voltage traces.

**Figure 5.**
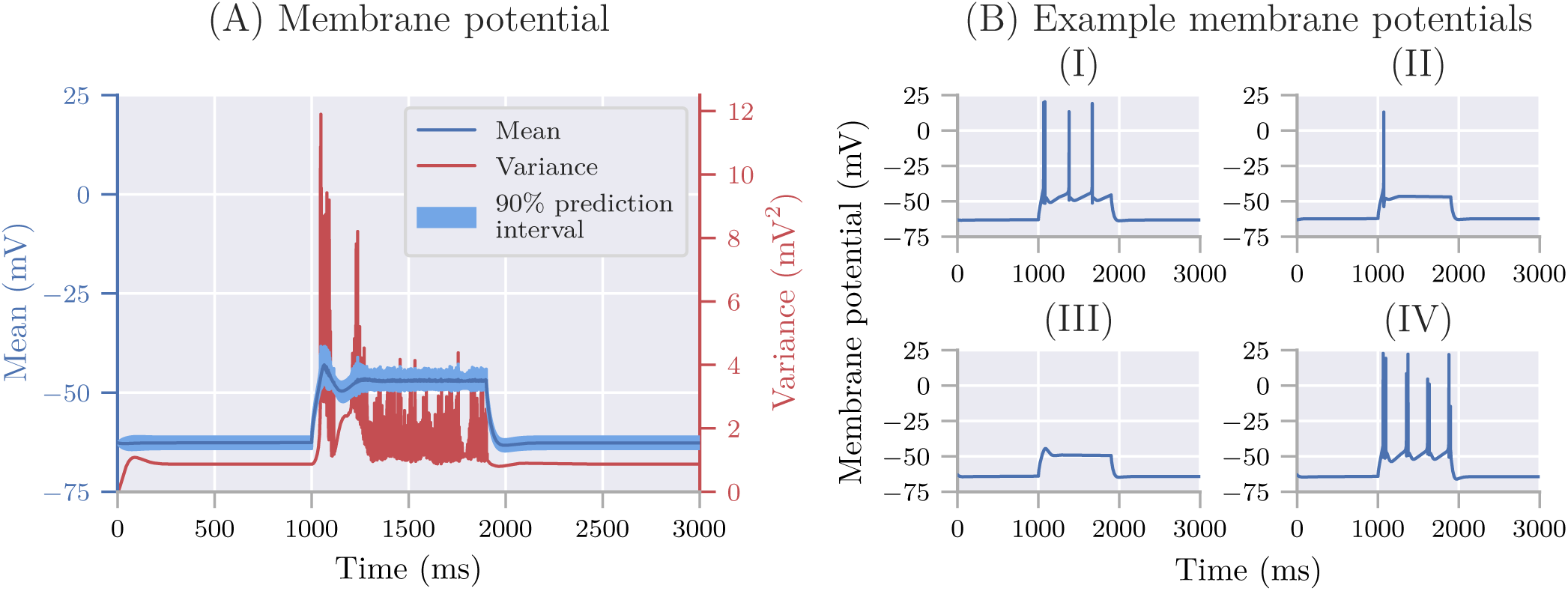
The uncertainty quantification and four selected model results for the interneuron model. (A) The mean and 90% prediction interval for the membrane potential of the interneuron model. (B) Four selected model outputs for different sets of parameters.

A feature-based analysis of the model output is shown in Figure 6. All statistical metrics from Section 2.2 and Section 2.3 are calculated by Uncertainpy for the model and each feature in `SpikingFeatures`, but for illustrative purposes we here only show the normalized sum of the first-order Sobol indices. See Section 3.4.3 for a description of each feature. As Figure 6 shows, different features are sensitive to different parameters. For example, the membrane potential of the neuron (panel C) is mainly sensitive to the reversal potential (*E*_pas_) of the passive current. The accommodation index (B) is most sensitive to the reversal potential. The spike rate (A), time before first spike (D), number of spikes (F), average action potential overshoot (G), and afterhyperpolarization depth (H) are most sensitive to the activation threshold of the Na^+^ channel (*Sh*_Na_). The average action potential overshoot (G) is also highly sensitive to the activation threshold of the K^+^ channel (*Sh*_Kdr_), as is the width of the action potential (E).

**Figure 6.**
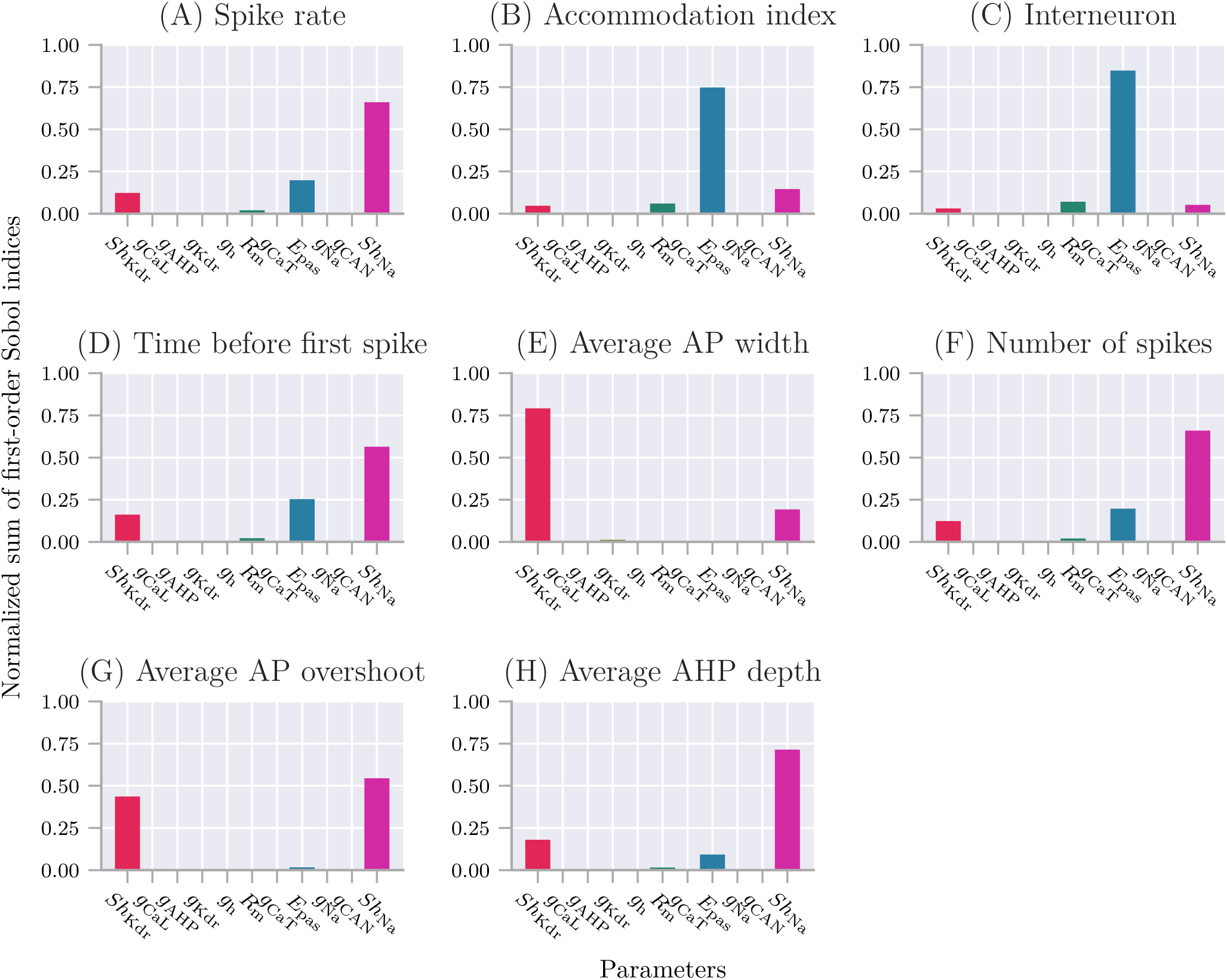
The sensitivity for features of the interneuron model. Normalized sum of the first-order Sobol indices for the thalamic interneuron model and features of the model. (A) Spike rate, that is, number of action potentials divided by stimulus duration. (B) Accommodation index, which is the normalized average difference in length of two consecutive interspike intervals. (C) Interneuron, which is the membrane potential of the model itself. (D) Time before first spike, that is, the time from stimulus onset to first elicited action potential. (E) Average AP width is the average action potential width taken at midpoint between the onset and peak of the action potential. (F) Number of spikes, which is the number of action potentials during stimulus period. (G) Average AP overshoot is the average action-potential peak voltage. (H) Average AHP depth, that is, the average minimum voltage between action potentials.

A feature-based sensitivity analysis like this gives valuable insight into the role of various biological mechanisms in determining the firing properties of a neuron. In the current model, the features in `SpikingFeatures` are almost exclusively sensitive to only four parameters (*E*_pas_, *R*_*m*_, *Sh*_Na_ and *Sh*_Kdr_), but the remaining parameters could be important for features not included in the relatively limited `SpikingFeatures` set. For example, one characteristic feature of thalamic interneurons is that they sometimes elicit characteristic bursts of action potentials (Zhu et al., 1999). Such a bursty behavior indeed occurs for some parameterizations of the model, as in Figure 5B (IV) where a closer inspection reveals that each of the four visible “thick spikes” actually consists of several spikes in rapid succession. The burstiness is likely to be sensitive to the T-type Ca^2+^ conductance (*g*_*CaT*_), which in previous modelling studies has been found important for burst generation (Zhu et al., 1999; Halnes et al., 2011; Allken et al., 2014). Several aspects of bursting activity is covered by `EfelFeatures`, and an analysis based on the extended feature list could likely reveal a higher sensitivity to *g*_*CaT*_ and possibly other model parameters. We do not here analyze additional features of the interneuron model, as the main purpose of the case study was to demonstrate the use of Uncertainpy on a detailed multicompartmental model.

### 4.4 Recurrent network of integrate-and-fire neurons

In the last case study, we use Uncertainpy to perform a feature-based analysis of the sparsely connected recurrent network of integrate-and-fire neurons by Brunel (2000). We implement the Brunel network using NEST inside a Python function, and create 10000 inhibitory and 2500 excitatory neurons. We record the output from 20 of the excitatory neurons, and simulate the network for 1000 ms. The code for this case study is found in `/uncertainpy/examples/brunel/`.

The Brunel model has four uncertain parameters, (i) the external input rate (*?*_ext_) relative to threshold rate (*v*_thr_) given as *η* = *v*_ext_/*v*_thr_, (ii) the relative strength of the inhibitory synapses *g*, (iii) the synaptic delay *D*, and (iv) the amplitude of excitatory postsynaptic current *J*_*e*_. Depending on the parameterizations of the model, the Brunel network may be in several different activity states. For the current case study we limit our analysis to two of these states, the synchronous regular (SR) state, where the neurons are almost completely synchronized, and the asynchronous irregular (AI) state, where the neurons fire individually at low rates. We create two sets of parameters and assume the parameter uncertainties are characterized by uniform probability distributions within the ranges shown in Table 4, which corresponds to the network being in each of the two states. Two selected model results representative of the network in both states are shown in Figure 7, which illustrates the difference between the two states. Figure 7 (A) shows the recorded spike trains for the Brunel network in the SR state between 200 ms and 300 ms of the simulation. This time window is representative of the network during the entire simulation after spiking has started, and is chosen to give more detailed overview due to the number of spikes that occur during the simulation. Figure 7 (B) shows the recorded spike trains for the Brunel network in the in the AI state for the entire simulation period.

**Table 4.** Parameters in the Brunel network for the asynchronous irregular (AI) and synchronous regular (SR) state. Each parameter has a uniform distribution within the given range.

| Parameter | Range SR | Range AI | Variable | Meaning |
| --- | --- | --- | --- | --- |
| $\eta$ | [1.5, 3.5] | [1.5, 3.5] | **eta** | External rate relative to threshold rate |
| $g$ | [1, 3] | [5, 8] | **g** | Relative strength of inhibitory synapses |
| $D$ | [1.5, 3] | [1.5, 3] | **delay** | Synaptic delay (ms) |
| $J_e$ | [0.05, 0.15] | [0.05, 0.15] | **J_e** | Amplitude excitatory postsynaptic current (mV) |

**Figure 7.**
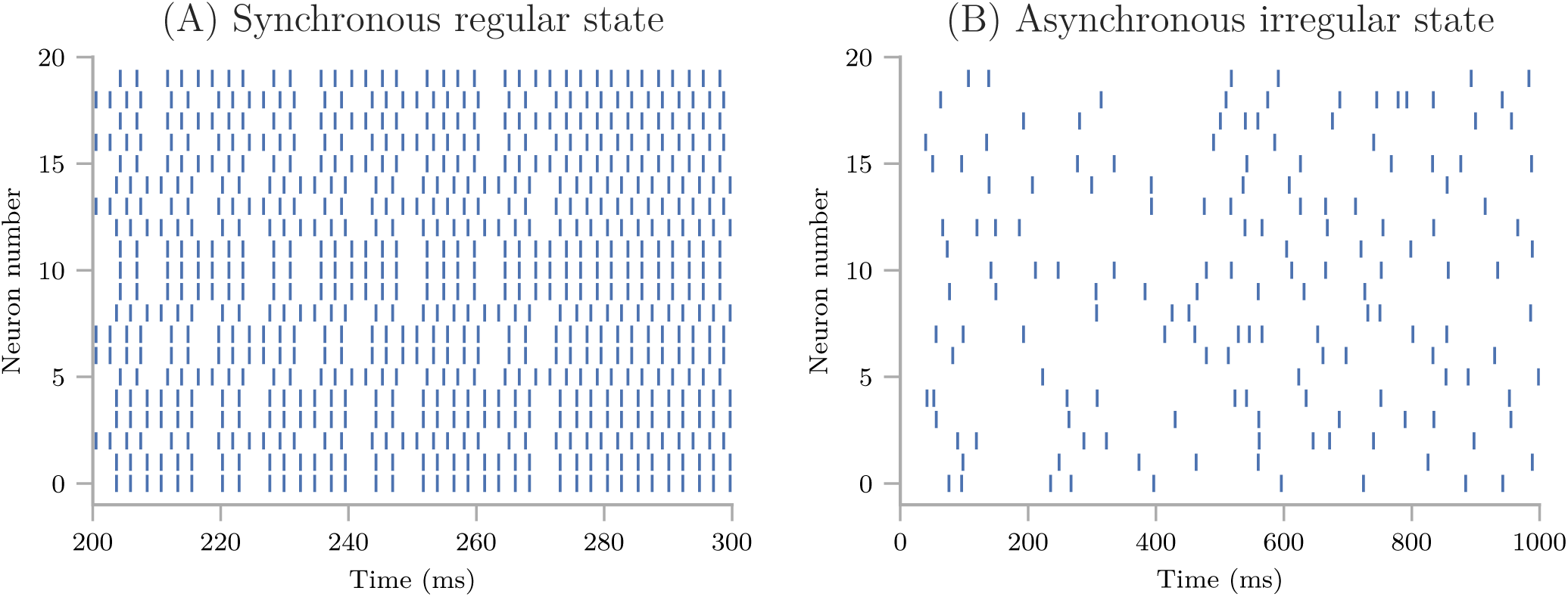
Example model results for the Brunel network. (A) The recorded spike train for the Brunel network in the synchronous regular state between 200 ms and 300 ms of the simulation. (B) The recorded spike trains for the Brunel network in the in the asynchronous irregular state for the entire simulation period. The network has 10000 inhibitory and 2500 excitatory neurons. We record the output from 20 of the excitatory neurons and simulate the network for 1000 ms.

We use the features in `NetworkFeatures` to examine features of the network dynamics. We perform an uncertainty quantification and sensitivity analysis for the model and all features for each of the network states, using polynomial chaos with point collocation. Of the 13 built-in network features in `NetworkFeatures`, we here only focus on two: the mean interspike interval and the correlation coefficient. These features are well suited to highlight the differences between the AI and SR network states.

#### 4.4.1 Mean interspike interval

The mean interspike interval is the average time it takes from a neuron elicits a spike until it elicits the next spike, averaged over all recorded neurons. The uncertainty quantification and sensitivity analysis for the mean interspike interval of the Brunel network are shown in Figure 8. The mean interspike interval is known to differ strongly between the SR and AI states, as found here. In the SR state, the mean of the mean interspike interval is low, with a comparable low variance reflecting the synchronous firing in the network. We can observe this in Figure 7A, where the interspike intervals are short and does not vary much. In the AI state, the mean of the mean interspike interval is high, with an even greater variance, reflecting the asynchronous firing in the network. This fits what we observe in Figure 7B, where the interspike intervals are relatively large and varies to a large degree.

**Figure 8.**
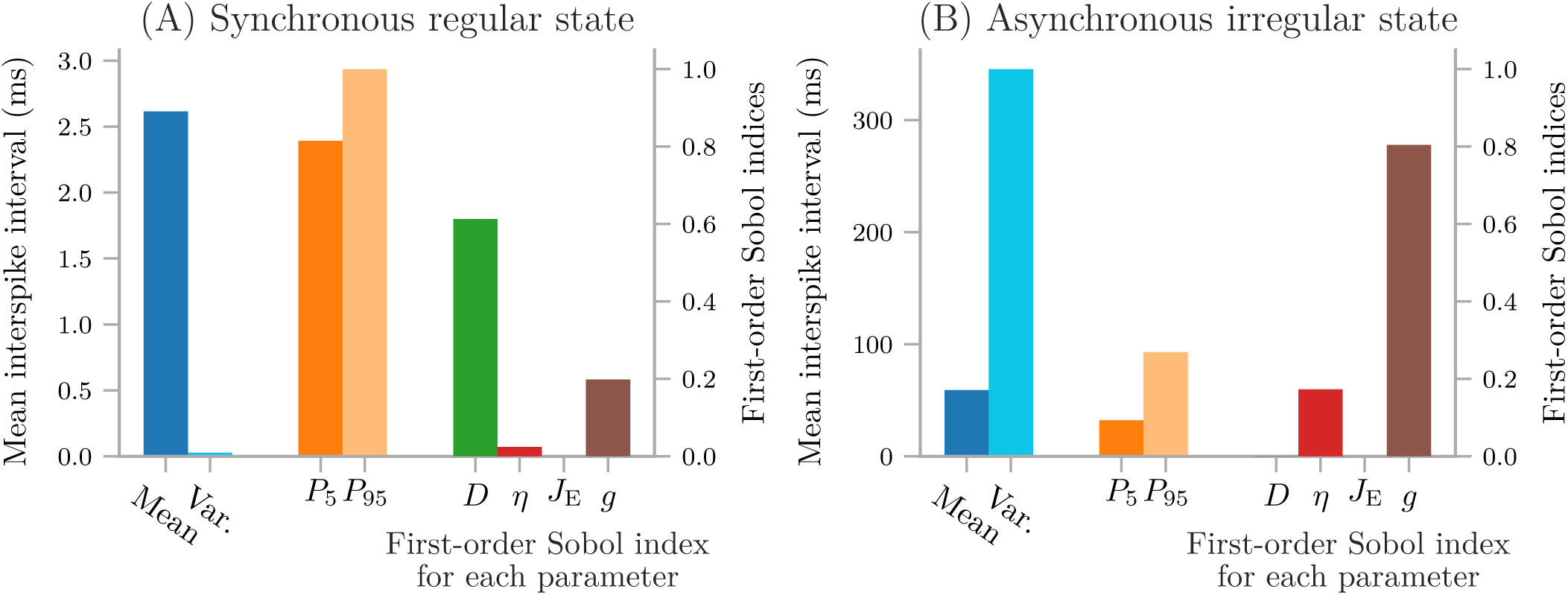
The mean interspike interval for the Brunel network in the two states. Mean, variance, 90% prediction interval, and first-order Sobol indices of the mean interspike interval of the Brunel network in the synchronous regular state (A), and in the asynchronous irregular state (B). The 90% prediction interval is indicated by the 5-th and 95-th percentiles, i.e., 90% of the mean spike intervals are between *P*_5_ and *P*_95_.

The two states were also found to be different in terms of which parameters the mean interspike interval is sensitive to. In the AI state, the network is almost exclusively sensitive to the relative strength of inhibitory synapses *g*, and only slightly sensitive to *η*, while in the SR state it is predominantly sensitive to the synaptic delay *D*, and less so to the relative synaptic strength *g*.

#### 4.4.2 Correlation coefficient

The pairwise Pearson’s correlation coefficient is a measure of how synchronous the spiking of a network is. This correlation coefficient measures the correlation between the spike trains of two neurons in the network. In Figure 9 we examine how the synchronicity in the Brunel network depends on parameter uncertainties by plotting the mean, variance, and first-order Sobol indices for the Pearson correlation coefficient in the SR and AI states.

**Figure 9.**
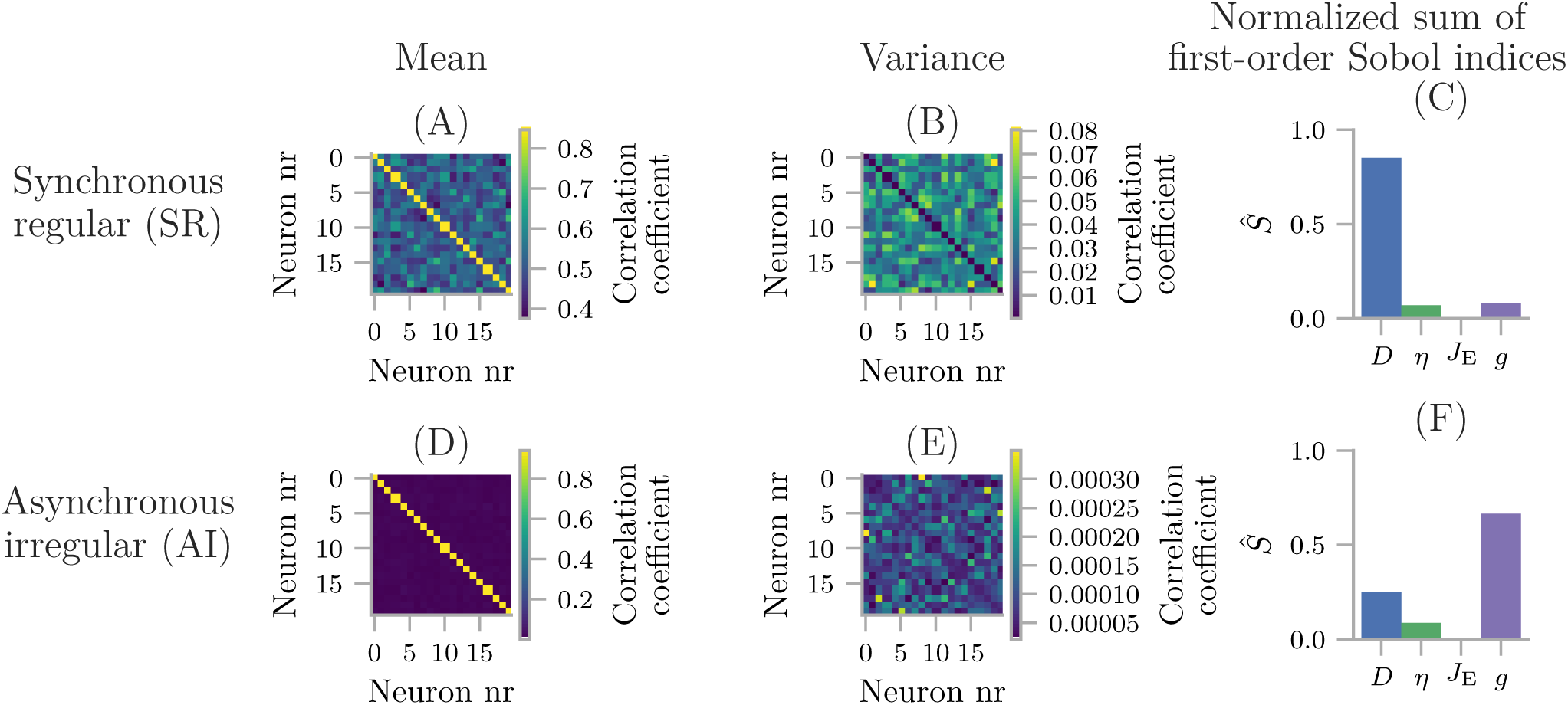
The pairwise Pearson’s correlation coefficient for the Brunel network in the two states. Mean (A, D), variance (B, E) and first-order Sobol indices (C, F) for the pairwise Pearson’s correlation coefficient of the Brunel network in the synchronous regular (A, B, C) and asynchronous irregular (D, E, F) states.

As expected from examining Figure 7, in the SR state (Figure 9A) the mean correlation coefficient between neurons is much higher than in the AI state (Figure 9D). The first-order Sobol indices further show that the degree of synchronicity is by far most sensitive to the synaptic delay *D* when the network is in the SR state (Figure 9C), and to the relative strength of inhibitory synapses *g* when the network is in the AI state (Figure 9F).

In terms of both features investigated here (the mean interspike interval and the pairwise Pearson’s correlation coefficient), the conclusions regarding model sensitivity are the same. The SR state of the Brunel network is most sensitive to the synaptic delay *D*, while the AI state is most sensitive to the relative strength of inhibitory synapses *g*. Neither of the states are particularly sensitive to the remaining parameters, that is the external input rate *η* and the excitatory postsynaptic current *J*_*E*_.

### 4.5 Additional examples

Additional examples for uncertainty quantification of the Izikevich neuron (Izhikevich, 2003), a reduced layer 5 pyramidal cell (Bahl et al., 2012), and a Hodgkin-Huxley model with shifted voltage (Sterratt et al., 2011) are found in `/uncertainpy/examples/`.

## DISCUSSION

A major challenge with models in neuroscience is that they tend to contain a number of uncertain parameters whose values are critical for the model behavior. In this paper we have presented Uncertainpy, a Python toolbox for uncertainty quantification and sensitivity analysis. Uncertainpy is tailored for neuroscience applications by its built-in capability for recognizing features in the model output.

The key aim of Uncertainpy is to make it quick and easy for the user to get started with uncertainty quantification and sensitivity analysis, without any need for detailed prior knowledge of uncertainty analysis. Uncertainpy is applicable to a range of different model types, as illustrated in the example applications. These included an uncertainty quantification and sensitivity analysis of four different models: a simple cooling coffee-cup model, the original Hodgkin-Huxley model for generation of action potentials, a NEURON model of a multicompartmental thalamic interneuron, and a NEST model of a sparsely connected recurrent network of integrate-and-fire neurons.

To our knowledge, Uncertainpy is the first toolbox to use polynomial chaos expansions to perform uncertainty quantification and sensitivity analysis in neuroscience. Compared to quasi-Monte Carlo methods, polynomial chaos expansions dramatically reduce the number of model evaluations needed to get reliable statistics. This is especially important for models that require a long simulation time, where uncertainty quantification using quasi-Monte Carlo methods could require an unfeasible amount of computer time.

### 5.1 Results of the case studies

The case studies examined in this paper were predominantly performed to demonstrate the use of Uncertainpy. We did not put much effort into estimating realistic distributions for the parameter uncertainties. For example, for the thalamic interneuron model (Section 4.3) we simply assumed that all parameters had uniform distributions within a*±*2.5% range of the values they had in the original model. This was a rather arbitrary choice, and it is unlikely to capture the real uncertainty distributions. Furthermore, it is likely that some parameters have broader uncertainty distributions than others. The conclusions from such an analysis should therefore be considered with caution, as the results depend on the chosen parameter distributions.

### 5.2 Applicability of Uncertainpy

It is not always evident which model is best suited to describe a certain system. For example, when we construct a neural model we first have to decide which mechanism (ion channels, ion pumps, synapses, etc.) to include in the model. Next we select a set of mathematical equations that describe these mechanisms. Such choices are seldom trivial, and no methods for resolving this structural uncertainty aspect of modeling are included in Uncertainpy. Uncertainpy focus on parameter uncertainties, which are important in their own right.

It is rarely the case that the distributions of the uncertain model parameters are precisely known. This implies that an exact quantification of the absolute output uncertainty can not be obtained. One way to deal with this is to limit the study to identify qualitative model predictions that are essentially parameter-independent (Bailey, 2001). However, despite a lack of precise knowledge of parameter uncertainties, quantitative measures such as those obtained with Uncertainpy may still give valuable insights in the relationship between model parameters and model output. This insight can guide experimentalists towards focusing on accurately measuring the parameters most critical for the model output. Additionally it can guide modellers by identifying mechanisms that can be sacrificed for model reduction purposes.

Obtaining the distributions of the uncertain parameters is mainly an empirical problem, and not a problem dealt with by Uncertainpy. The uncertainties of a parameter can generally be divided into two main classes: aleatory uncertainties and epistemic uncertainties. Epistemic uncertainty reflects a lack of knowledge, and can in principle be reduced to zero by acquiring additional information. Measurement uncertainty is one example of epistemic uncertainty. Aleatory uncertainty, on the other hand, is uncertainty due to inherent variability of the parameters. Causes of aleatory uncertainty are biological variability (Edelman and Gally, 2001) or regulation (Marder and Goaillard, 2006). The importance of distinguishing between aleatory and epistemic uncertainties has evoked some debate (Hora, 1996; Ferson and Ginzburg, 1996; Oberkampf et al., 2002; Ferson et al., 2004; Kiureghian and Ditlevsen, 2009; Mullins et al., 2016), but the distinction is at least important for how to interpret the results of an uncertainty quantification. Due to inherent variability, the parameters do not have true fixed values, but rather distributions of possible values. The variance in the model output is in such cases an expression of biological variation. On the other hand, parameters with epistemic uncertainties produce an uncertainty as to whether or not we have gotten “the correct result”.

### 5.3 Further development of Uncertainpy

There are several ways that Uncertainpy can be further developed, and we list some of these in this section. Currently, Uncertainpy only works with Python 2 due to limitations in some of the packages utilized, so one obvious improvement is to implement support for Python 3.

If a model or features of a model uses adaptive time steps, Uncertainpy performs an interpolation of the output to get the regular result needed in the uncertainty quantification and sensitivity analysis. Currently, Uncertainpy only has support for interpolation of one-dimensional output (vectors). Depending on requests from the community, this aspect can be improved.

The current version of Uncertainpy can only perform sensitivity analysis when using polynomial chaos expansions. This is the best choice of method for models with relatively few parameters. However, polynomial chaos expansions scales worse with the number of uncertain parameters than quasi-Monte Carlo methods, and the latter may be the best choice for models with many (typically > 20) parameters. To make Uncertainpy more flexible in terms of model size, a natural extension would be to include methods for performing sensitivity analysis based on quasi-Monte Carlo methods.

Another possibility is to implement a screening method, such as the Morris One-At-a-Time method (Morris, 1991), to be able to use polynomial chaos expansions on models with many parameters. The idea of the screening is to reduce the number of uncertain parameters by setting the parameters that have the least effect on the model output to fixed values. We can then use only the parameters with the greatest effect on the model output when performing the uncertainty quantification and sensitivity analysis.

The built-in feature library in Uncertainpy can easily be expanded by adding additional features. The number of built-in simulators can also easily be extended. We encourage the users to add custom features and models through Github pull requests.

### 5.4 Outlook

In many fields of the physical sciences, the model parameters that goes into simulations are known with high accuracy. For example, in quantum mechanical simulations of molecular systems, the masses of the nuclei and the electrons, as well as the parameters describing their electrical interaction, are known so precisely that uncertainty in model parameters is not an issue (Marx and Hutter, 2009). This is not the case in computational biology in general, and in computational neuroscience in particular. Model parameters of biological systems often have an inherent variability, and some may even be actively regulated and change with time. They can therefore not be precisely known. We therefore consider uncertainty quantification and sensitivity analysis to be particularly important in biology.

Uncertainpy was developed with the aim of enabling such analysis, by providing a tool for precise evaluation of the effect of uncertain model parameters on the model predictions, which makes it easy and quick to get started with uncertainty analysis, without any need for detailed prior knowledge. Being an open-source Python toolbox, we hope that Uncertainpy can be further developed through a joint effort within the neuroscience community.

## AUTHOR CONTRIBUTIONS

ST, GH, and GE conceived of and designed the project. ST designed, wrote, tested and documented the software and performed analysis of the examples. ST, GH, and GE wrote and revised the paper.

## ACKNOWLEDGMENTS

We would like to acknowledge the vital contribution to the vision of this work by Hans-Petter Langtangen, who regretfully passed away before he could see the work complete. We would also acknowledge the help from Jonathan Feinberg in teaching the basis of polynomial chaos expansions, as well as how to use Chaospy.

## Footnotes

1 https://github.com/simetenn/uncertainpy

2 https://github.com/simetenn/uncertainpy

3 http://efel.readthedocs.io

## References

Achard, P. and De Schutter, E. (2006). Complex parameter landscape for a complex neuron model. PLoS Computational Biology 2, 0794–0804. doi:10.1371/journal.pcbi.0020094

Allken, V., Chepkoech, J.-L., Einevoll, G. T., and Halnes, G. (2014). The subcellular distribution of T-type Ca2+ channels in interneurons of the lateral geniculate nucleus. PloS one 9, e107780. doi:10.1371/journal.pone.0107780

Archer, G. E. B., Saltelli, A., and Sobol, I. M. (1997). Sensitivity measures,anova-like Techniques and the use of bootstrap. Journal of Statistical Computation and Simulation 58, 99–120. doi:10.1080/00949659708811825

Babtie, A. C. and Stumpf, M. P. H. (2017). How to deal with parameters for whole-cell modelling. Journal of The Royal Society Interface 14. doi:10.1098/rsif.2017.0237

Bahl, A., Stemmler, M., Herz, A., and Roth, A. (2012). Automated optimization of a reduced layer 5 pyramidal cell model based on experimental data. Journal of Neuroscience Methods 210, 22–34. doi:10.1016/j.jneumeth.2012.04.006

Bailey, J. E. (2001). Complex biology with no parameters. Nature Biotechnology 19, 503. doi:doi:10.1038/89204

Beck, M. B. (1987). Water quality modeling: A review of the analysis of uncertainty. Water Resources Research 23, 1393–1442. doi:10.1029/WR023i008p01393

Beer, R. D., Chiel, H. J., and Gallagher, J. C. (1999). Evolution and analysis of model CPGs for walking: II. General principles and individual variability. Journal of computational neuroscience 7, 119–47. doi:10.1023/a:1008920021246

Bhalla, U. S. and Bower, J. M. (1993). Exploring parameter space in detailed single neuron models: simulations of the mitral and granule cells of the olfactory bulb. J Neurophysiol 69, 1948–1965

Blomquist, P., Devor, A., Indahl, U. G., Ulbert, I., Einevoll, G. T., and Dale, A. M. (2009). Estimation of thalamocortical and intracortical network models from joint thalamic single-electrode and cortical laminar-electrode recordings in the rat barrel system. PLOS Computational Biology 5, 1–24. doi:10.1371/journal.pcbi.1000328

Blot, A. and Barbour, B. (2014). Ultra-rapid axon-axon ephaptic inhibition of cerebellar Purkinje cells by the pinceau. Nature Neuroscience 17, 289–295. doi:10.1038/nn.3624

Blue Brain Project (2015). efel. Available online at: https://github.com/BlueBrain/eFEL (Accessed October 12, 2017). Commit: d252bc12c7dec3f47c55f6ea8a1f8110a80a18aa

Borgonovo, E. and Plischke, E. (2016). Sensitivity analysis: A review of recent advances. European Journal of Operational Research 248, 869–887. doi:10.1016/j.ejor.2015.06.032

Brodland, G. W. (2015). How computational models can help unlock biological systems. Seminars in Cell and Developmental Biology 47-48, 62–73. doi:10.1016/j.semcdb.2015.07. 001

Brunel, N. (2000). Dynamics of sparsely connected networks of excitatory and inhibitory spiking neurons. Journal of Computational Neuroscience 8, 183–208. doi:10.1023/A: 1008925309027

Collette, A. (2013). Python and HDF5 (O’Reilly)

Crestaux, T.,;tre, O., and L e MaîMartinez, J. M. (2009). Polynomial chaos expansion for sensitivity analysis. Reliability Engineering and System Safety 94, 1161–1172. doi:10.1016/ j.ress.2008.10.008

Dayan, P. and Abbott, L. F. (2001). Theoretical Neuroscience: Computational and Mathematical Modeling of Neural Systems (The MIT Press)

De Schutter, E. and Bower, J. M. (1994). An active membrane model of the cerebellar Purkinje cell II. Simulation of synaptic responses. Journal of neurophysiology 71, 401–419. doi:10.1152/jn.1994.71.1.401

Degenring, D., Froemel, C., Dikta, G., and Takors, R. (2004). Sensitivity analysis for the reduction of complex metabolism models. Journal of Process Control 14, 729 – 745. doi:https://doi.org/10.1016/j.jprocont.2003.12.008. Dynamics, Monitoring, Control and Optimization of Biological Systems

Druckmann, S., Banitt, Y., Gidon, A. A., Schürmann, F., Markram, H., and Segev, I. (2007). A novel multiple objective optimization framework for constraining conductance-based neuron models by experimental data. Frontiers in Neuroscience 1, 7–18. doi:10.3389/neuro.01.1.1.001.2007

Eck, V. G., Donders, W. P., Sturdy, J., Feinberg, J., Delhaas, T., Hellevik, L. R., et al. (2016). A guide to uncertainty quantification and sensitivity analysis for cardiovascular applications. International Journal for Numerical Methods in Biomedical Engineering 32, e02755–n/a. doi:10.1002/cnm.2755

Edelman, G. M. and Gally, J. A. (2001). Degeneracy and complexity in biological systems. Proceedings of the National Academy of Sciences 98, 13763–13768. doi:10.1073/pnas. 231499798

Einevoll, G. T. (2009). Sharing with Python. Frontiers in Neuroscience 3, 334–335. doi:10. 3389/neuro.01.037.2009

Eppler, J. M., Pauli, R., Peyser, A., Ippen, T., Morrison, A., Senk, J., et al. (2015). Nest 2.8.0. doi:10.5281/zenodo.32969

Feinberg, J. and Langtangen, H. P. (2015). Chaospy: An open source tool for designing methods of uncertainty quantification. Journal of Computational Science 11, 46–57. doi:10. 1016/j.jocs.2015.08.008

Ferson, S. and Ginzburg, L. R. (1996). Different methods are needed to propagate ignorance and variability. Reliability Engineering and System Safety 54, 133–144. doi:10.1016/ S0951-8320(96)00071-3

Ferson, S., Joslyn, C. A., Helton, J. C., Oberkampf, W. L., and Sentz, K. (2004). Summary from the epistemic uncertainty workshop: Consensus amid diversity. Reliability Engineering and System Safety 85, 355–369. doi:10.1016/j.ress.2004.03.023

Friedrich, P., Vella, M., Gulyás, A. I., Freund, T. F., and Káli, S. (2014). A flexible, interactive software tool for fitting the parameters of neuronal models. Frontiers in Neuroinformatics 8, 1–19. doi:10.3389/fninf.2014.00063

Garcia, S., Guarino, D., Jaillet, F., Jennings, T., Pröpper, R., Rautenberg, P. L., et al. (2014). Neo: an object model for handling electrophysiology data in multiple formats. Frontiers in Neuroinformatics 8, 1–10. doi:10.3389/fninf.2014.00010

Glen, G. and Isaacs, K. (2012). Estimating Sobol sensitivity indices using correlations. Environmental Modelling and Software 37, 157–166. doi:10.1016/j.envsoft.2012.03.014

Goldman, M. S., Golowasch, J., Marder, E., and Abbott, L. F. (2001). Global structure, robustness, and modulation of neuronal models. The Journal of neuroscience: the official journal of the Society for Neuroscience 21, 5229–38. doi:21/14/5229[pii]

Golowasch, J., Goldman, M. S., Abbott, L. F., and Marder, E. (2002). Failure of averaging in the construction of a conductance-based neuron model. Journal of neurophysiology 87, 1129–1131. doi:10.1152/jn.00412.2001

Gutenkunst, R. N., Waterfall, J. J., Casey, F. P., Brown, K. S., Myers, C. R., and Sethna, J. P. (2007). Universally sloppy parameter sensitivities in systems biology models. PLoS Computational Biology 3, 1871–1878. doi:10.1371/journal.pcbi.0030189

Halnes, G., Augustinaite, S., Heggelund, P., Einevoll, G. T., and Migliore, M. (2011). A multi-compartment model for interneurons in the dorsal lateral geniculate nucleus. PLoS Computational Biology 7, 1–12. doi:10.1371/journal.pcbi.1002160

Halnes, G., Liljenström, H., and Århem, P. (2007). Density dependent neurodynamics. BioSystems 89, 126–134

Halnes, G., Ulfhielm, E., Eklöf Ljunggren, E., Kotaleski, J. H., and Rospars, J. P. (2009). Modelling and sensitivity analysis of the reactions involving receptor, G-protein and effector in vertebrate olfactory receptor neurons. Journal of Computational Neuroscience 27, 471–491. doi:10.1007/s10827-009-0162-6

Hamby, D. M. (1994). A review of techniques for parameter sensitivity analysis of environmental models. Environmental Monitoring and Assessment 32, 135–154. doi:10. 1007/BF00547132

Hammersley, J. M. (1960). Monte carlo methods for solving multivariable problems. Annals of the New York Academy of Sciences 86, 844–874. doi:10.1111/j.1749-6632.1960.tb42846. x

Hay, E., Hill, S., Schürmann, F., Markram, H., and Segev, I. (2011). Models of neocortical layer 5b pyramidal cells capturing a wide range of dendritic and perisomatic active properties. PLoS Computational Biology 7. doi:10.1371/journal.pcbi.1002107

Hay, E., Schürmann, F., Markram, H., and Segev, I. (2013). Preserving axosomatic spiking features despite diverse dendritic morphology. Journal of neurophysiology 109, 2972–81. doi:10.1152/jn.00048.2013

Hines, M. L. and Carnevale, N. T. (1997). The NEURON Simulation Environment. Neural Computation 9, 1179–1209. doi:10.1162/neco.1997.9.6.1179

Hodgkin, A. L. and Huxley, A. F. (1952). A quantitative description of membrane current and its application to conduction and excitation in nerve. J Physiol 117, 500–544. doi:10. 1016/S0092-8240(05)80004-7

Homma, T. and Saltelli, A. (1996). Importance measures in global sensitivity analysis of nonlinear models. Reliability Engineering & System Safety 52, 1–17. doi:10.1016/ 0951-8320(96)00002-6

Hora, S. C. (1996). Aleatory and epistemic uncertainty in probability elicitation with an example from hazardous waste management. Reliability Engineering and System Safety 54, 217–223. doi:10.1016/S0951-8320(96)00077-4

Hosder, S., Walters, R., and Balch, M. (2007). Efficient Sampling for Non-Intrusive Polynomial Chaos Applications with Multiple Uncertain Input Variables. 48th AIAA/ASME/ASCE/AHS/ASC Structures, Structural Dynamics, and Materials Conference doi:10.2514/6.2007-1939

Izhikevich, E. M. (2003). Simple model of spiking neurons 14, 1569–1572. doi:10.1109/TNN. 2003.820440

Izhikevich, E. M. and Edelman, G. M. (2008). Large-scale model of mammalian thalamocortical systems. Proceedings of the National Academy of Sciences 105, 3593–3598. doi:10.1073/pnas.0712231105

Kiureghian, A. D. and Ditlevsen, O. (2009). Aleatory or epistemic? Does it matter? Structural Safety 31, 105–112. doi:10.1016/j.strusafe.2008.06.020

Koch, C. and Segev, I. (eds.) (1998). Methods in Neuronal Modeling: From Ions to Networks (MIT Press), 2nd edn.

Kuchibhotla, K. V., Gill, J. V., Lindsay, G. W., Papadoyannis, E. S., Field, R. E., Sten, T. A., et al. (2017). Parallel processing by cortical inhibition enables context-dependent behavior. Nature Neuroscience 20, 62–71. doi:10.1038/nn.4436

Leamer, E. (1985). Sensitivity Analyses Would Help. Amer 75, 308–313

Lemieux, C. (2009). *Monte Carlo and Quasi-Monte Carlo Sampling*. Springer Series in Statistics (Dordrecht: Springer). doi:10.1007/978-0-387-78165-5

Mäki-Marttunen, T., Halnes, G., Devor, A., Metzner, C., Dale, A. M., Andreassen, O. A., et al. (2018). A stepwise neuron model fitting procedure designed for recordings with high spatial resolution: Application to layer 5 pyramidal cells. Journal of Neuroscience Methods 293, 264–283. doi:10.1016/j.jneumeth.2017.10.007

Marder, E. and Goaillard, J. M. (2006). Variability, compensation and homeostasis in neuron and network function. Nature Reviews Neuroscience 7, 563–574. doi:10.1038/nrn1949

Marder, E. and Taylor, A. L. (2011). Multiple models to capture the variability in biological neurons and networks. Nature Neuroscience 14, 133–138. doi:10.1038/nn.2735

Marino, S., Hogue, I. B., Ray, C. J., and Kirschner, D. E. (2008). A methodology for performing global uncertainty and sensitivity analysis in systems biology. Journal of Theoretical Biology 254, 178–196. doi:10.1016/j.jtbi.2008.04.011

Markram, H., Muller, E., Ramaswamy, S., Reimann, M. W., Abdellah, M., Sanchez, C. A., et al. (2015). Reconstruction and Simulation of Neocortical Microcircuitry. Cell 163, 456–492. doi:10.1016/j.cell.2015.09.029

Marx, D. and Hutter, J. (2009). Ab initio Molecular Dynamics: Basic Theory and Advanced Method (Cambridge University Press)

McKerns, M. M., Strand, L., Sullivan, T., Fang, A., and Aivazis, M. A. G. (2012). Building a Framework for Predictive Science, 1–12

Merolla, P. A., Arthur, J. V., Alvarez-Icaza, R., Cassidy, A. S., Sawada, J., Akopyan, F., et al. (2014). A million spiking-neuron integrated circuit with a scalable communication network and interface. Science 345, 668–673. doi:10.1126/science.1254642

Morris, M. D. (1991). Factorial sampling plans for preliminary computational experiments. Technometrics 33, 161–174

Muller, E., Bednar, J. A., Diesmann, M., Gewaltig, M.-O., Hines, M., and Davison, A. P. (2015). Python in neuroscience. Frontiers in Neuroinformatics 9, 14–17. doi:10.3389/ fninf.2015.00011

Mullins, J., Ling, Y., Mahadevan, S., Sun, L., and Strachan, A. (2016). Separation of aleatory and epistemic uncertainty in probabilistic model validation. Reliability Engineering and System Safety 147, 49–59. doi:10.1016/j.ress.2015.10.003

Najm, H. N. (2009). Uncertainty Quantification and Polynomial Chaos Techniques in Computational Fluid Dynamics. Annual Review of Fluid Mechanics 41, 35–52. doi:10. 1146/annurev.fluid.010908.165248

Narayan, A. and Jakeman, J. (2014). Adaptive Leja sparse grid constructions for stochastic collocation and high-dimensional approximation doi:10.1137/140966368

NeuralEnsemble (2017). Elephant - electrophysiology analysis toolkit. Available online at: https://github.com/NeuralEnsemble/elephant (Accessed November 27, 2017). Commit: f04e966857fd047243fb736f72df60d10f65ff79

Oberkampf, W. L., DeLand, S. M., Rutherford, B. M., Diegert, K. V., and Alvin, K. F. (2002). Error and uncertainty in modeling and simulation. Reliability Engineering and System Safety 75, 333–357. doi:10.1016/S0951-8320(01)00120-X

O’Donnell, C., Gonçalves, J. T., Portera-Cailliau, C., and Sejnowski, T. J. (2017). Beyond excitation/inhibition imbalance in multidimensional models of neural circuit changes in brain disorders. eLife 6, e26724. doi:10.7554/eLife.26724

Oliphant, T. E. (2007). Python for scientific computing. Computing in Science Engineering 9, 10–20. doi:10.1109/MCSE.2007.58

Pozzorini, C., Mensi, S., Hagens, O., Naud, R., Koch, C., and Gerstner, W. (2015). Automated High-Throughput Characterization of Single Neurons by Means of Simplified Spiking Models. PLoS Computational Biology 11, 1–29. doi:10.1371/journal.pcbi.1004275

Prinz, A. A., Bucher, D., and Marder, E. (2004). Similar network activity from disparate circuit parameters. Nature Neuroscience 7, 1345–1352. doi:10.1038/nn1352

Rifkin, R. M. and Lippert, R. A. (2007). Notes on regularized least squares. Massachusetts Institute of Technology

Rosenblatt, M. (1952). Remarks on a Multivariate Transformation. The Annals of Mathematical Statistics 23, 470–472. doi:10.1214/aoms/1177729394

Rossa, A., Liechti, K., Zappa, M., Bruen, M., Germann, U., Haase, G., et al. (2011). The COST 731 Action: A review on uncertainty propagation in advanced hydro-meteorological forecast systems. Atmospheric Research 100, 150–167. doi:10.1016/j.atmosres.2010.11.016

Rubinstein, R. Y. and Kroese, D. P. (2016). Simulation and the Monte Carlo Method (John Wiley & Sons, Inc.). doi:10.1002/9781118631980

Saltelli, A. (2002). Sensitivity analysis for importance assessment 22, 579–590. doi:10.1111/ 0272-4332.00040

Saltelli, A., Annoni, P., Azzini, I., Campolongo, F., Ratto, M., and Tarantola, S. (2010). Variance based sensitivity analysis of model output. Design and estimator for the total sensitivity index. Computer Physics Communications 181, 259–270. doi:10.1016/j.cpc. 2009.09.018

Schulz, D. J., Goaillard, J.-M., and Marder, E. E. (2007). Quantitative expression profiling of identified neurons reveals cell-specific constraints on highly variable levels of gene expression. Proceedings of the National Academy of Sciences 104, 13187–13191. doi:10.1073/pnas.0705827104

Smolyak, S. (1963). Quadrature and interpolation formulas for tensor products of certain classes of functions. Dokl. Akad. Nauk SSSR 148, 1042–1045

Snowden, T. J., van der Graaf, P. H., and Tindall, M. J. (2017). Methods of model reduction for large-scale biological systems: A survey of current methods and trends. Bulletin of Mathematical Biology 79, 1449–1486. doi:10.1007/s11538-017-0277-2

Sobol, I. M. (1990). Sensitivity analysis for nonlinear mathematical models. Matematicheskoe Modelirovanie 2, 112–118. doi:10.18287/0134-2452-2015-39-4-459-461.

Sterratt, D., Graham, B., Gillies, A., and Willshaw, D. (2011). Principles of Computational Modelling in Neuroscience (Cambridge University Press). doi:10.1017/ CBO9780511975899

Sudret, B. (2008). Global sensitivity analysis using polynomial chaos expansions. Reliability Engineering and System Safety 93, 964–979. doi:10.1016/j.ress.2007.04.002

Svensson, C. M., Coombes, S., and Peirce, J. W. (2012). Using evolutionary algorithms for fitting high-dimensional models to neuronal data. Neuroinformatics 10, 199–218. doi:10. 1007/s12021-012-9140-7

Taylor, A. L., Goaillard, J.-M., and Marder, E. (2009). How multiple conductances determine electrophysiological properties in a multicompartment model. J Neurosci. doi:10.1523/ JNEUROSCI.4438-08.2009

Tobin, A.-E. (2006). Endogenous and Half-Center Bursting in Morphologically Inspired Models of Leech Heart Interneurons. Journal of Neurophysiology 96, 2089–2106. doi:10. 1152/jn.00025.2006

Torres Valderrama, A., Witteveen, J., Navarro, M., and Blom, J. (2015). Uncertainty Propagation in Nerve Impulses Through the Action Potential Mechanism. Journal of mathematical neuroscience 5, 3. doi:10.1186/2190-8567-5-3

Turanyi, T. and Turányi, T. (1990). Sensitivity Analysis of Comprex Kinetic Systems. Tools and Applications. Journal of Mathematical Chemistry 5, 203–248. doi:10.1007/bf01166355

Van Geit, W., Achard, P., and De Schutter, E. (2007). Neurofitter: a parameter tuning package for a wide range of electrophysiological neuron models. Frontiers in neuroinformatics 1, 1. doi:10.3389/neuro.11.001.2007

Van Geit, W., De Schutter, E., and Achard, P. (2008). Automated neuron model optimization techniques: A review. Biological Cybernetics 99, 241–251. doi:10.1007/ s00422-008-0257-6

Van Geit, W., Gevaert, M., Chindemi, G., Rössert, C., Courcol, J.-D., Muller, E. B., et al. (2016). BluePyOpt: Leveraging Open Source Software and Cloud Infrastructure to Optimise Model Parameters in Neuroscience. Frontiers in Neuroinformatics 10, 1–18. doi:10.3389/fninf.2016.00017

Vanier, M. C. and Bower, J. M. (1999). A comparative survey of automated parameter-search methods for compartmental neural models. Journal of Computational Neuroscience 7, 149–171. doi:10.1023/A:1008972005316

Wang, H. and Sheen, D. A. (2015). Combustion kinetic model uncertainty quantification, propagation and minimization. Progress in Energy and Combustion Science 47, 1–31. doi:10.1016/j.pecs.2014.10.002

Wood-Schultz, D. H. S. &. M. M. (2011). QMU and Nuclear Weapons Certification-What’s under the Hood? Los Alamos Science 44, 55–82

Xiu, D. (2010). Numerical methods for stochastic computations: A spectral method approach (Princeton, NJ, USA: Princeton University Press)

Xiu, D. and Hesthaven, J. S. (2005). High-Order Collocation Methods for Differential Equations with Random Inputs. SIAM Journal on Scientific Computing 27, 1118–1139. doi:10.1137/040615201

Yildirim, B. and Karniadakis, G. E. (2015). Stochastic simulations of ocean waves: An uncertainty quantification study. Ocean Modelling 86, 15–35. doi:10.1016/j.ocemod.2014. 12.001

Zhu, J. J., Uhlrich, D. J., and Lytton, W. W. (1999). Burst firing in identified rat geniculate interneurons. Neuroscience 91, 1445–1460

Zi, Z. (2011). Sensitivity analysis approaches applied to systems biology models. IET Systems Biology 5, 336–346(10)

